# Phage proteins block and trigger retron toxin/antitoxin systems

**DOI:** 10.1101/2020.06.22.160242

**Authors:** Jacob Bobonis, Karin Mitosch, André Mateus, George Kritikos, Johanna R. Elfenbein, Mikhail M. Savitski, Helene Andrews-Polymenis, Athanasios Typas

## Abstract

Bacteria carry dozens of Toxin/Antitoxin systems (TAs) in their chromosomes. Upon growth, the antitoxin is co-expressed and neutralizes the toxin. TAs can be activated and inhibit growth, but when and how this occurs has largely remained enigmatic, hindering our understanding of their physiological roles. We developed TIC/TAC (Toxin Inhibition/Activation Conjugation), a high-throughput reverse genetics approach, to systematically identify molecular blockers and triggers of TAs. By applying TIC/TAC to a tripartite TA, the retron-Sen2 of *Salmonella* Typhimurium, we have identified multiple blockers and triggers of phage origin. We demonstrate that diverse phage functionalities are sensed by the DNA-part of the antitoxin and ultimately activate the retron toxin. Phage-origin proteins can circumvent activation by directly blocking the toxin. Some identified triggers and blockers also act on an *E. coli* retron-TA, Eco9. We propose that retron-TAs act as abortive-infection anti-phage defense systems, and delineate mechanistic principles by which phages trigger or block them.

## INTRODUCTION

Toxin/Antitoxin systems (TA) are prokaryotic bipartite operons consisting of an antitoxin and toxin gene pair. The antitoxin encodes a protein or an RNA, which counteracts the protein toxin. The first such system was discovered in *E. coli*, where *ccdB*/*ccdA* “addicts” cells in stably inheriting the F-plasmid ^1^. Plasmid-based TAs lead to addiction, because the growth of cells losing the plasmid is inhibited by the toxin, which becomes free as the antitoxin is more labile ^2,3^. TAs confer addictive phenotypes by additional mechanisms and/or in other mobile elements ^4,5^. The genomics revolution unraveled that TA systems are also ubiquitous in bacterial chromosomes ^6^. For instance, *Escherichia coli* K-12 encodes at least 35 chromosomal-TA systems ^7^, whereas there are more than 80 TA systems of a specific type (type II) in the chromosome of *Mycobacterium tuberculosis* H37Rv ^8,9^. TA systems are frequently found in the accessory genomes of bacteria and change drastically between strains of the same species ^10^. Yet, for the vast majority, we have currently little insight into what triggers them, and hence their physiological role remains elusive and intensely debated ^11^.

A handful of TA systems have been shown to protect bacteria against phages via abortive infection (Abi) ^12–17^. Abi is a general term describing bacterial defenses activated after a phage bypasses the bacterial innate or adaptive antiviral systems (such as Restriction-Modification ^18^ or CRISPR-Cas systems ^19^). In contrast to innate/adaptive systems which directly target the phage, Abi systems inhibit the growth of the infected bacterium, indirectly reducing phage progeny at the population-level, at the expense of the infected cell ^20^. Although the molecular triggers of Abi systems are unknown, it is presumed that such TAs can somehow sense phage infection and activate their toxin. Interestingly, a T4-phage protein (Dmd) has been found to directly bind, and inhibit the chromosomal toxins of two such TA systems, thereby enabling the T4 phage to infect a TA-containing *E. coli* ^21,22^. Thus, phages carry genes to counteract toxins of Abi TA systems (blocker genes). We reasoned that the identification of triggers or blockers of TA systems can propel our understanding of their role in bacteria.

In the accompanying manuscript, we report that the *Salmonella enterica* retron-Sen2 (historically named retron-ST85 ^23^) encodes a novel tripartite TA system. The toxin RcaT is directly counteracted by an antitoxin unit formed by the reverse transcriptase (RT) bound to a multi-copy single-stranded DNA (msDNA) ^24^. To elucidate its physiological role, we have developed a reverse genetics approach that enables the systematic discovery of TA blocker and trigger genes. TIC/TAC (Toxin Inhibition/Activation Conjugation) uses plasmid libraries to survey the role of all possible genome-encoded molecular cues, and takes advantage of the two-sided phenotype associated with all TAs: ectopically inducing expression of a toxin inhibits bacterial growth, while co-expressing it with its antitoxin restores growth. Using two genome-wide *E. coli* overexpression libraries, we identified dozens of triggers and blockers for the retron-Sen2, enriched in phage-origin proteins. Phage proteins are sensed by the retron-TA via multiple mechanisms, which are inherent to the tripartite architecture of the TA system. We propose that the retron-TA acts as an anti-phage defense system, and we provide a method that can be readily applied to uncover the physiological role of any TA system.

## RESULTS

### A new systematic method for identifying TA blockers and triggers

To survey for molecular cues of TA systems, we used genome-wide *E. coli* single-gene overexpression libraries (MOB; p1 ^25^ and TransBac; p2 ^26^) in tandem with strains carrying appropriate TA-expressing plasmids (Fig. 1A). We reasoned that this would allow us to identify genes that trigger and block the TA system, but are normally silent or buffered when cells are growing in lab conditions. We used a tripartite retron-TA system ^24^ from *Salmonella enterica* subsp. enterica ser. Typhimurium str. 14028s (*S*Tm) as the input TA system. The retron-TA (retron-Sen2 ^27^) encodes the toxin RcaT, which is inhibited directly by a complex formed between the msDNA and the reverse transcriptase (msDNA-RT) ^24^. Overexpressing *rcaT* causes toxicity in *E. coli*, while overexpressing the entire retron-TA (*msrmsd*-*rcaT*-*rrtT*) alleviates the RcaT-mediated toxicity ^24^.

**Figure 1.**
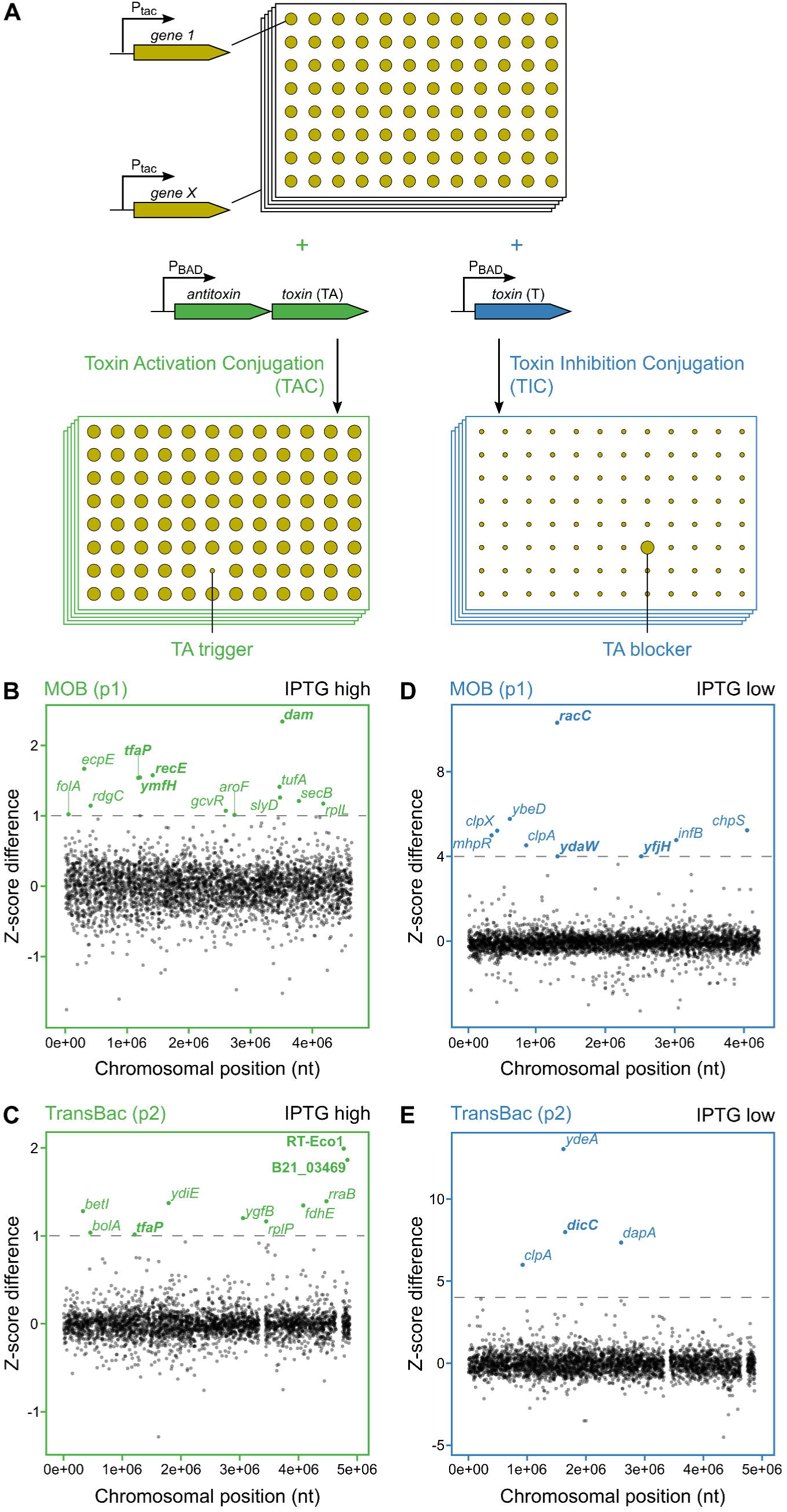
Toxin Inhibition/Activation Conjugation (TIC/TAC): a high-throughput reverse genetics approach to discover TA blockers and triggers. **(A)** Two systematic, arrayed, conjugative-proficient *E. coli* gene overexpression libraries (Ptac-gene 1 to Ptac-gene X; library-plasmids ^25,26^) are transferred into recipient strains of the desired genetic background, which carry compatible P_BAD_-controlled plasmids, expressing either an antitoxin-toxin (P_BAD_-TA), or a toxin (P_BAD_-T). Inducing the P_BAD_-T plasmid inhibits growth, while inducing the P_BAD_-TA plasmid does not. Co-inducing library-plasmids carrying a TA-trigger results into TA-mediated growth inhibition (TAC), while co-inducing library-plasmids carrying a TA-blocker alleviates toxin-mediated growth inhibition (TIC). **(B-C)** TAC screen using the IPTG-inducible library-plasmids from the MOB (B; p1) ^25^ and TransBac (C; p2) ^26^ libraries, respectively. Arrayed libraries (384-density format) were conjugated in an *E. coli* recipient carrying an arabinose-inducible p-retron plasmid. Transconjugants carrying both plasmids were selected by antibiotics and auxotrophies (see Methods), individual colony integral opacities were measured ^65^, and z-scores of strains were calculated after growing on experiment plates containing arabinose + low (0.1 mM) or high (1 mM) IPTG, or on control plates containing only low or high IPTG. Z-score differences per strain (y-axes) were derived by subtracting z-scores in control plates, from z-scores in experiment plates (*n = 2*; see Table S2 for scores). Scores shown from high-IPTG induction (see ED Fig. 1A-B for low-IPTG results). Chromosomal position (x-axes) is according to the gene start and *E. coli* MG1655 coordinates. Grey line denotes the hit cut-off (Z-score difference >1), and phage-related trigger-genes are in bold. **(D-E)** TIC screen using the IPTG-inducible library-plasmids from the MOB (**D**; p1) ^25^ and TransBac (**E**; p2) ^26^ libraries, respectively. Conjugation was performed as in **B-C**, but with an *E. coli* recipient carrying an arabinose-inducible p-*rcaT* plasmid. Axes and data analysis as described in **B-C**, but z-score differences were derived by subtracting z-scores of experiment plates from control plates. Scores shown from low-IPTG induction (see ED Fig. 3A for MOB high-IPTG results). Grey line denotes the hit cut-off (Z-score difference >4), and phage-related blocker-genes are in bold.

To identify retron-TA triggers and blockers we developed two reverse genetics fitness-based screens called TAC (Toxin Activation Conjugation) and TIC (Toxin Inhibition Conjugation), respectively (Fig. 1A). Wildtype *E. coli* BW25113 carrying either an arabinose-inducible p-retron plasmid (P_BAD_-*msrmsd*-*rcaT-rrtT*; TAC) or p-toxin plasmid (P_BAD_-*rcaT*) ^24^ was mated with the two *E. coli* overexpression libraries (Ptac-*gene-X*), by high-throughput conjugation on agar plates. After selection, the resulting 18,434 double-plasmid bearing strains (9,217 for each screen) were grown under different induction conditions: co-inducing the retron/toxin and the library-plasmid (see TIC/TAC procedure in Methods).

We reasoned that strains that could grow upon library-plasmid or p-retron induction, but were growth-inhibited upon co-induction, carried library-plasmids containing a retron-TA trigger that activates RcaT (Fig. 1A; TAC). To identify these strains, we compared the fitness of every individual strain between the library-plasmid induction condition (IPTG; control plates) and the corresponding co-inducing conditions (Arabinose + IPTG; experiment plates) (see TIC/TAC analysis in Methods). We identified 13 triggers from the MOB library (Fig. 1B) and 10 triggers from the TransBac library (Fig. 1C) upon high trigger-induction. Fewer and slightly different triggers were identified by lower IPTG induction (ED Fig. 1A-B). Despite the high reproducibility of individual screens (ED Fig. 1C), the overlap in hits between the two libraries was low (ED Fig. 1D). This inconsistency has many reasons: stringent cutoffs (hence false negatives), quality of libraries (missing genes, plasmid mutations, or cloning errors), and importance of trigger levels (e.g., MOB vectors are medium-copy ^25^, while TransBac vectors are single-copy ^26^).

To ensure that the identified triggers inhibit growth by specifically activating RcaT, we selected 15 triggers, sequenced the plasmid-inserts, and conjugated them again into *E. coli* strains expressing either an empty vector, p-retron, p-retron-Δ*rcaT*, or p-*rcaT*. All 15 were benign when co-expressed only with the antitoxin (p-retron-Δ*rcaT* plasmid), but inhibited growth when expressed with the full retron (ED Fig. 2A), suggesting that they trigger RcaT. As in the screen, different triggers displayed varying degrees of RcaT-activation, and required different IPTG induction-levels to manifest their effect (ED Fig. 2B). We noticed that several triggers, especially strong ones, were prophage-encoded genes (enrichment p-val = 0.01-0.025, depending on library and induction level – Table S1) –*recE* (Rac-prophage), *tfaP* & *ymfH* (e14-prophage), retron RT-Eco1 (P2-like-prophage gene in *E. coli* BL21), and B21_03469 (prophage in *E. coli* BL21). Although *dam* is part of the *E. coli* core genome, Dam methylases are also commonly found in phages ^28^. Other triggers belonged to house-keeping processes, such as translation (*tufA, rplL, rplP, rplV)*, DNA-binding/transcription (*rdgC, rraB, betI, bolA*), protein quality control and translocation (*slyD, secB, yajC*), chorismate-tetrahydrofolate biosynthesis (*folA, gcvR, aroF, aroK*), but also orphan genes (*ygfB, ydiE, ecpE, gnsA*).

**Figure 2.**
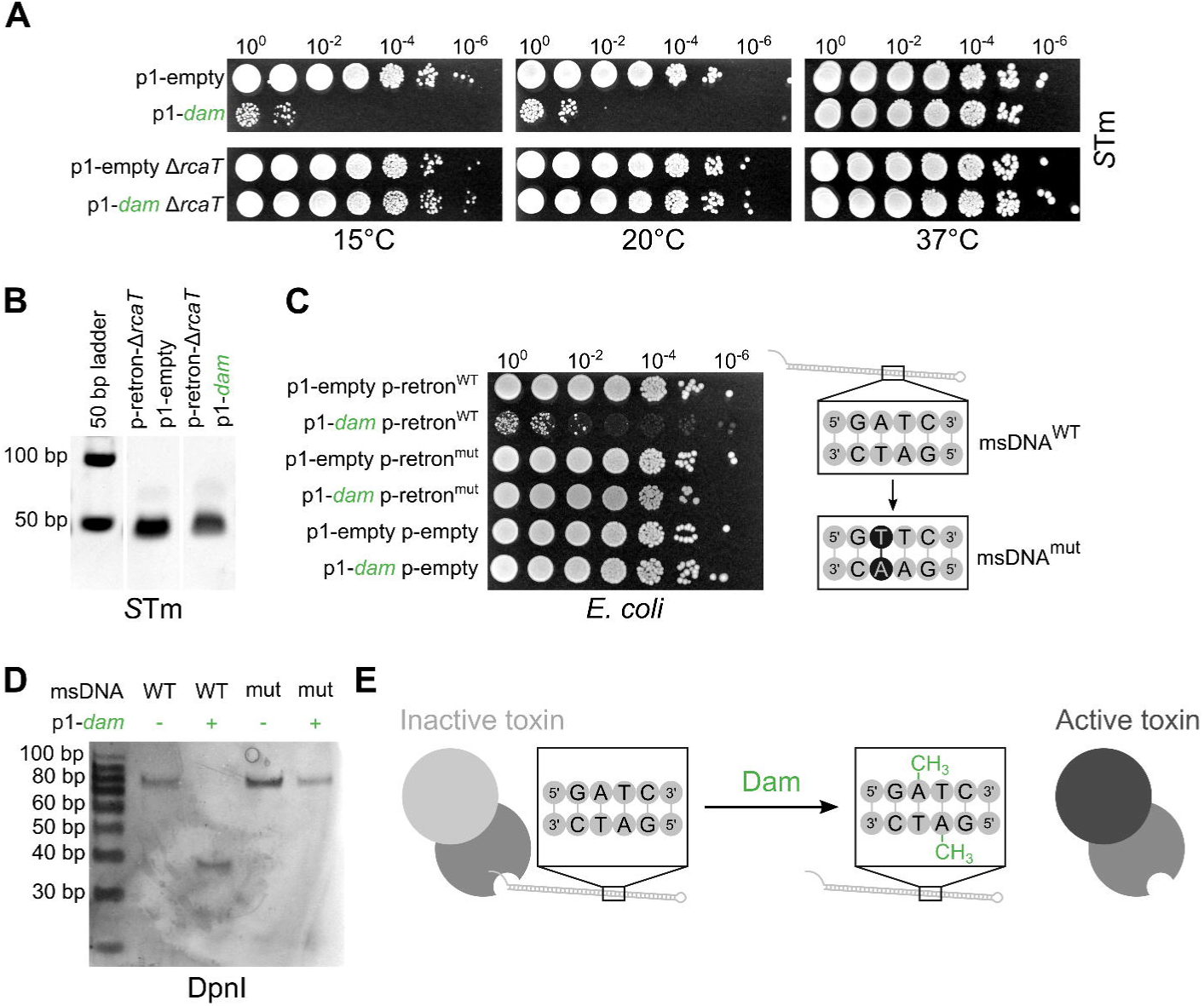
Dam triggers the retron-TA by directly methylating msDNA. **(A)** Overexpressing Dam triggers the endogenous *S*Tm retron-TA. *S*Tm strains (WT, Δ*rcaT*) carrying plasmid p1-*dam* or an empty vector (p1-empty) were grown for 5 hours in ampicillin-LB, serially diluted, spotted on ampicillin-LB plates containing IPTG (low), and then incubated at 15°C, 20°C, or 37°C. Representative data shown from two independent experiments. **(B)** Dam overexpression does not impact msDNA levels. msDNA were isolated from *S*Tm strains carrying combinations of plasmids p-retron-Δ*rcaT*, p1-*dam*, or p1-empty. Plasmids were co-induced with arabinose and IPTG (low). Extracted msDNA was electrophoresed in a TBE-Polyacrylamide gel. A representative gel of three independent experiments is shown. **(C)** Mutating the 5’-GATC-3’ motif in the msDNA duplex abolishes Dam-triggering, without affecting duplex formation. *E. coli* BW25113 carrying combinations of plasmids p-retron^WT^, p-retron^mut^ (5’-GTTC-3’ mutation), p1-*dam*, and empty vectors (p-empty, p1-empty) were grown for 5 hours in LB with appropriate antibiotics, serially diluted, spotted on LB plates containing antibiotics, IPTG (low), and arabinose, and incubated at 37°C. Representative data shown from two independent experiments. **(D)** Dam methylates the 5’-GATC-3’ site on msDNA. msDNA was isolated from *S*Tm strains carrying plasmids p-retron^WT^ or p-retron^mut^, and msDNA^WT^/msDNA^mut^ were purified further by gel-extraction. Purified msDNA was digested with DpnI, which recognizes and cuts methylated 5’-GATC-3’ sites, and digests were electrophoresed on a denaturing TBE-polyacrylamide gel (msDNA runs higher in denaturing compared to native polyacrylamide gels). **(E)** Dam triggers the retron-TA by methylating the msDNA. This results into an active RcaT (active toxin in black) – here shown to occur by dissociation of the msDNA from the RT (hypothetical).

Conversely to the TAC screen, we reasoned that strains able to grow upon p-*rcaT* induction, carried library plasmids containing toxin blockers that inhibited RcaT, and thereby allowing for growth (Fig. 1A; TIC). We identified 9 blockers from the MOB (Fig. 1D) and 4 blockers from the TransBac library (Fig. 1E) upon low blocker induction. As for the TAC screen, data were reproducible, but most blockers were specific to library (except for *clpA*) and induction conditions (ED Fig. 3). To validate these hits, we selected 11 blockers, sequenced their plasmid-inserts, and conjugated them back into *E. coli* strains expressing either a p-empty, a p-*rcaT*, a p-retron, or a p-retron-Δ*rcaT* plasmid. All blockers specifically alleviated RcaT-toxicity, while the overall fitness remained unchanged (ED Fig. 4A). Blockers inhibited RcaT to a different degree, depending on IPTG induction levels (ED Fig. 4B). Blockers were also enriched in prophage genes (enrichment p-val < 0.01 for MOB library and aggregated results – Table S1) – *racC, ydaW* (Rac-prophage), *yfjH* (CP4-57-prophage), *yjhC* (KpLE2-prophage), and *dicC* (Qin-prophage). Other blockers were related to protein quality control (*clpA, clpX, dnaJ*), translation (*infB, dtd, tcdA*), TA systems (*chpS*), conditionally-induced pathways (*mhpR, mhpE, ybeD, dsrB, glpE, ugpC*) and genes affecting mating or induction levels (*dapA* & *ydeA*). In summary, we developed a systematic approach to identify TA blockers and triggers (TIC/TAC), which we used to study the tripartite TA system, retron-Sen2. Many triggers and blockers had phage origins, prompting us to investigate further their underlying links to the retron.

**Figure 3.**
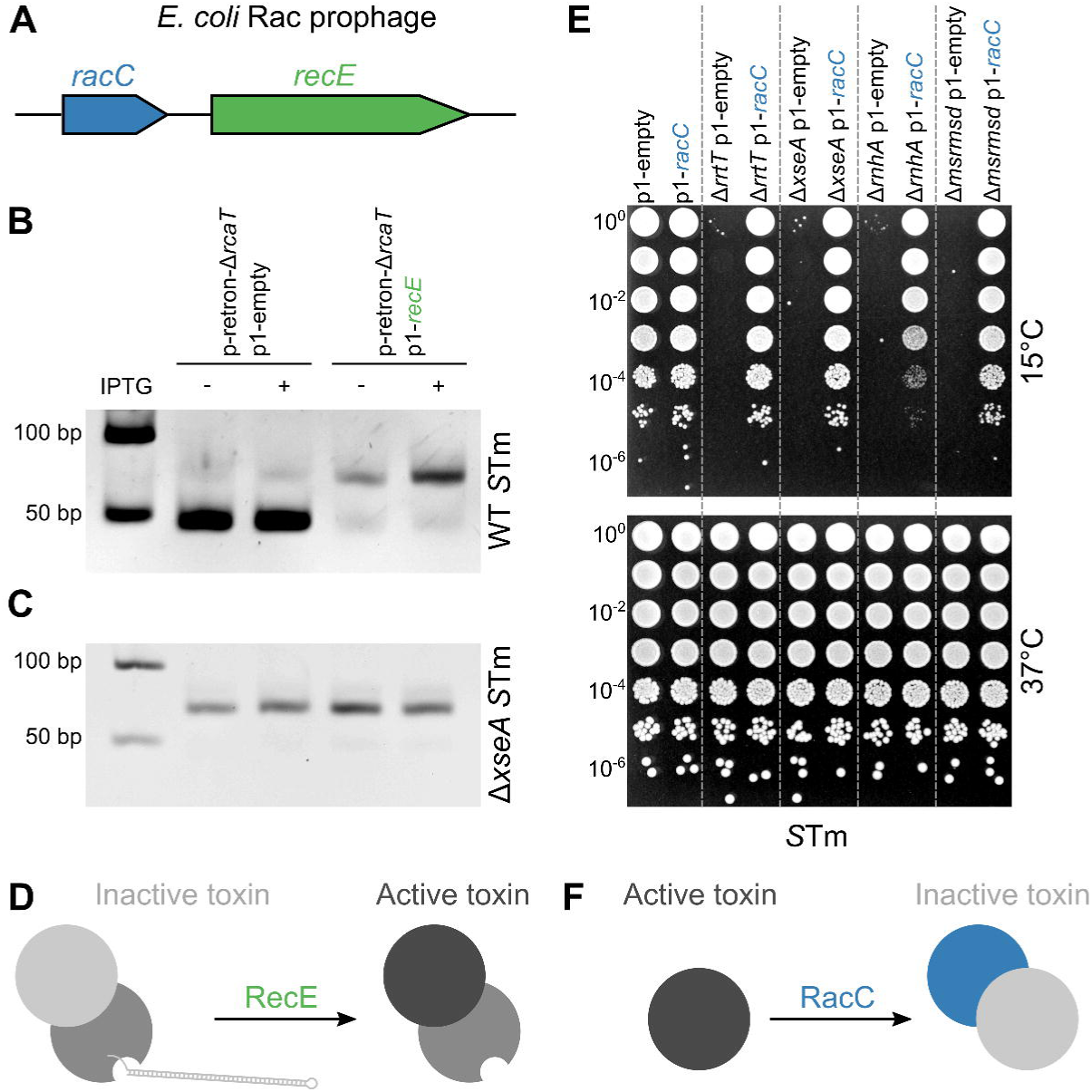
Rac prophage genes *racC*-*recE* are a blocker-trigger gene-pair. **(A)** *racC* and *recE* are linked Rac-prophage *E. coli* genes. The Rac prophage in *E. coli* BW25113 contains *recE* (exodeoxyribonuclease VIII), preceded by a small orphan gene, *racC*. **(B)** RecE degrades mature msDNA *in vivo*. msDNA was extracted from *S*Tm strains carrying combinations of plasmids p-retron-Δ*rcaT* with p1-*recE*, or p1-empty, respectively. Plasmids were co-induced with arabinose and IPTG (low). Extracted msDNA were electrophoresed in a TBE-Polyacrylamide gel. Mature msDNA runs close to the 50bp ladder band (double stranded DNA), whereas unprocessed DNA runs slightly higher and accumulates upon RecE overexpression. Representative results from three independent experiments are shown. **(C)** Immature msDNA cannot be degraded by RecE *in vivo*. Same as in panel **B**, but msDNA were isolated from *S*Tm Δ*xseA* strains. Representative results from two independent experiments are shown. **(D)** Model of RecE triggering the retron-TA by degrading msDNA. RecE directly degrades the mature msDNA, resulting in dissociation of the RT-msDNA complex, and activation of RcaT. **(E)** RacC blocks cold-sensitivity of endogenous *S*Tm retron-antitoxin deletions. *S*Tm strains (WT, Δ*rrtT*, Δ*xseA*, Δ*rnhA*, Δ*msrmsd*) carrying plasmid p1-*racC* or an empty vector (p1-empty) were grown for 5 hours in ampicillin-LB, serially diluted, spotted on ampicillin-LB plates containing IPTG (low), and incubated at 15°C, or 37°C. Representative data shown from two independent experiments. **(F)** Model of RacC blocking the retron-TA by inhibiting RcaT. RacC inhibits the toxic activity of RcaT, presumably by directly binding to RcaT.

**Figure 4.**
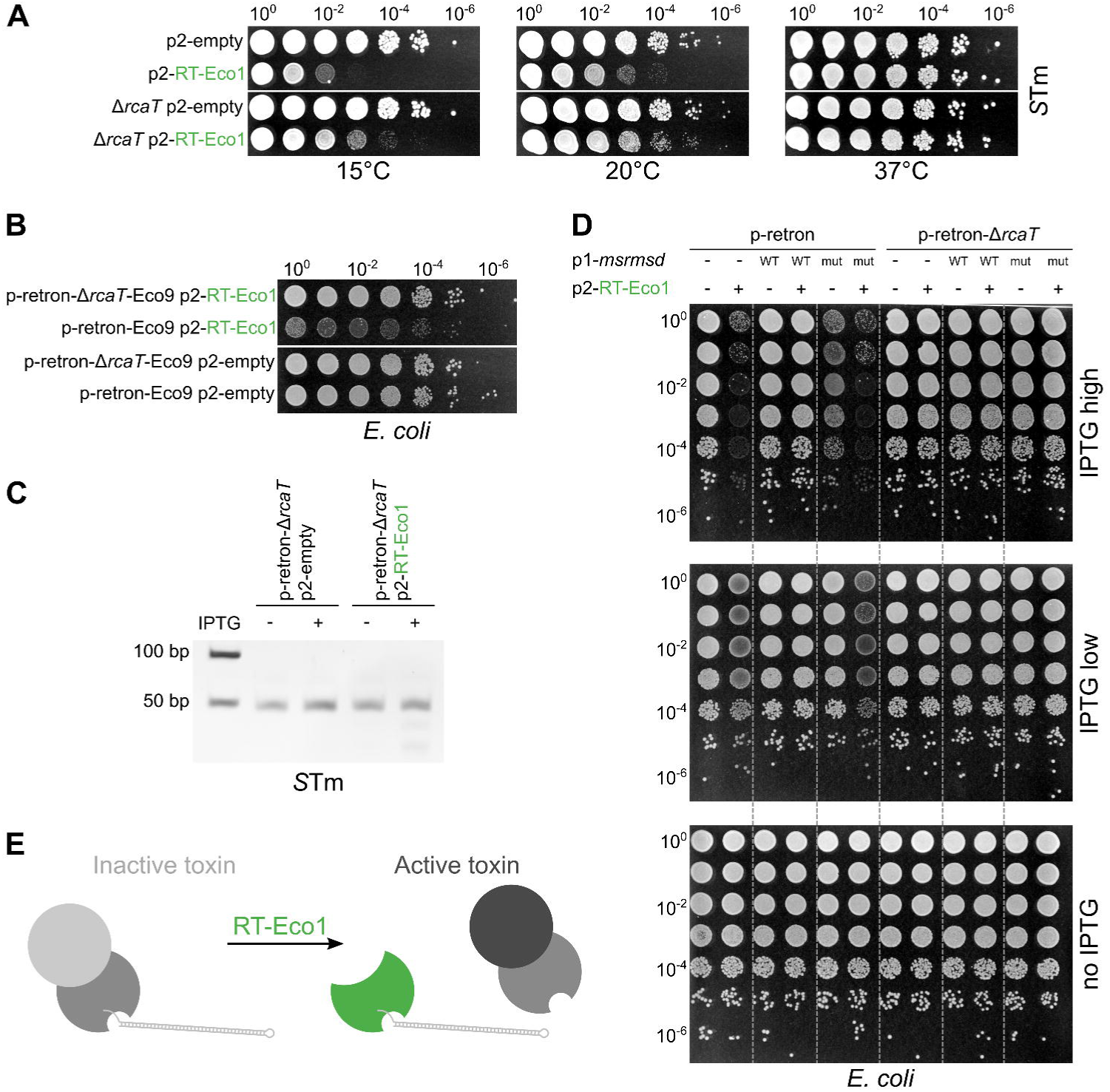
RT-Eco1 triggers the retron by sequestering the msDNA-Sen2 from RT-Sen2. **(A)** Overexpressing RT-Eco1 partially triggers the endogenous *S*Tm retron-TA. *S*Tm strains (WT, Δ*rcaT*) carrying plasmid p2-RT-Eco1 or an empty vector (p2-empty) were grown for 5 hours in tetracycline-LB, serially diluted, spotted on tetracycline-LB plates containing IPTG (low), and incubated at 15°C, 20°C, or 37°C. Representative data shown from two independent experiments. **(B)** RT-Eco1 triggers the retron-Eco9 from *E. coli* NILS-16 ^24^. *E. coli* BW25113 carrying combinations of plasmids p-retron-Eco9, p-retron-Δ*rcaT*-Eco9, p2-RT-Eco1, and p2-empty, were grown for 5 hours in LB with appropriate antibiotics, serially diluted, spotted on LB plates with antibiotics, arabinose, and IPTG (high), and incubated at 37°C. Representative data shown from three independent experiments. **(C)** RT-Eco1 does not affect msDNA levels. msDNA was extracted from *S*Tm Δretron cells being complemented by p-retron-Δ*rcaT*, and carrying p2-RT-Eco1 or p2-empty. Plasmids were co-induced with arabinose and IPTG (high). Extracted msDNAs were electrophoresed in a TBE-Polyacrylamide gel, and a representative gel from two independent experiments is shown. **(D)** msDNA overexpression alleviates RT-Eco1 triggering. *E. coli* BW25113 carrying combinations of plasmids p-retron, p-retron-Δ*rcaT*, p1-*msrmsd*^WT^, p1-*msrmsd*^mut^, p2-RT-Eco1, and empty vectors (p1-empty, p2-empty) were grown and spotted as in **B**, but in different antibiotics and IPTG concentrations. **(E)** RT-Eco1 triggers the retron-TA by sequestering msDNA away from its native RT. RT-Eco1 titrates msDNA-Sen2, resulting in unloaded RT-Sen2, which cannot alone neutralize the toxicity of RcaT.

### Dam triggers the retron-TA by directly methylating the antitoxin msDNA

In *E. coli*, the primary DNA adenine methylase is Dam ^29^, which methylates adenines in 5’-GATC-3’ DNA duplexes ^30^. Dam was the sole retron-TA trigger that activated RcaT-toxicity even without IPTG-induction (leaky levels; ED Fig. 2). Homologues of Dam with the same methylation recognition-site are present in other bacteria (e.g., *S*Tm ^31^), as well as in phages (e.g., phage P1 ^32^). Plasmids expressing *dam* from *S*Tm, or from phage P1, also triggered the retron-TA (ED Fig. 5A), suggesting that the 5’-GATC-3’ adenine methylation activity itself could be causing the retron-TA triggering.

**Figure 5.**
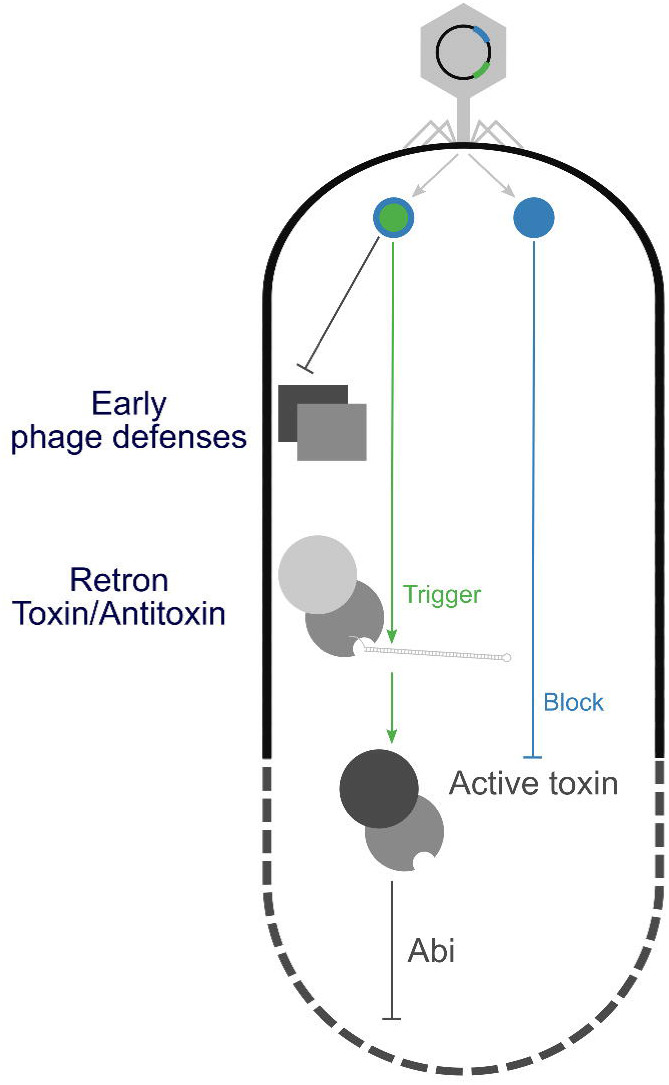
Retrons are Abi systems protecting against phage defense. Phages encode anti-restriction proteins (green-blue circle) to counteract bacterial Restriction-Modification systems (early-defense systems; RM). These anti-RM proteins (Dam, RecE) trigger the retron-TA by directly interacting with the msDNA, which disrupts the antitoxin. The latter allows the toxin (RcaT) to inhibit the growth of the infected bacterium, and thereby to indirectly stop the phage propagation – abortive infection (Abi). Thus, phage proteins are simultaneously blockers for bacterial innate defenses and triggers of secondary Abi defense systems (hence green-blue color). Being an arms race, phages have also evolved blocker proteins (blue), able to directly inhibit toxin activity of triggered Abi systems (e.g., RacC).

Upon perturbations in msDNA biosynthesis, the endogenous RcaT in *S*Tm inhibits growth in cold-temperatures ^24^. Overexpressing *dam* in *S*Tm phenocopied this, inhibiting growth in a *rcaT*-dependent manner at 15°C and 20°C, but not at 37°C (Fig. 2A). This finding implied that Dam may act by directly inactivating the antitoxin unit (RT-msDNA). To test if this occurs by reducing the msDNA levels, we isolated msDNA from strains carrying a *dam* overexpression plasmid, or an empty vector. msDNA levels were similar in the two conditions (Fig. 2B), suggesting that *dam* overexpression does not affect msDNA levels.

Although msDNAs are single-stranded ^27^, they are all reverse-transcribed from msd-RNAs containing inverted-repeats, thereby ultimately forming extended DNA hairpins, i.e., double-stranded DNA (dsDNA) ^33^. Dam methylates adenines in 5’-GATC-3’ in dsDNA. We noticed that the msDNA produced by the retron-TA contained this motif in its hairpin. To test whether msDNA methylation is involved in Dam-mediated retron-TA triggering, we mutated the Dam recognition motif on the msDNA, while retaining intact the hairpin structure (5’-GTTC-3’ duplex; p-retron^mut^). While co-expressing *dam* with the wildtype msDNA inhibited growth (5’-GATC-3’ duplex; p-retron^WT^), co-expressing *dam* with the p-retron^mut^ resulted in no change in fitness (Fig. 2C). Importantly, the p-retron^mut^ did not affect the RcaT toxicity, since expression of p-retron^mut^ impaired growth in a Δ*xseA E. coli*, which produces an immature msDNA that cannot support a functional antitoxin unit ^24,34^ (ED Fig. 5B). Furthermore, the p-retron^mut^ did not affect the levels or stability of msDNA (ED Fig. 5C). Overall, these results strongly suggested that Dam triggers RcaT by methylating a specific adenine of the 5’-GATC-3’ duplex on msDNA.

To prove that the msDNA did get methylated, we took advantage of the specificity of the restriction enzyme DpnI, which cleaves only adenine-methylated 5’-GATC-3’ duplexes ^35^. We purified wildtype (5’-GATC-3’) or mutated msDNA (5’-GTTC-3’) from strains carrying a *dam* or an empty plasmid, and digested them with DpnI. DpnI could only cut wildtype msDNA derived from a *dam* overexpressing strain, but not msDNA isolated from a strain with an empty vector, nor mutated msDNA (Fig. 2D). This also suggests that the msDNA is insufficiently methylated by the endogenous Dam methylase (all strains still carry *dam* in chromosome), which is consistent with endogenous Dam-levels being rate limiting for higher-copy DNA elements ^36^. Thus, Dam methylates the msDNA hairpin at the specific 5’-GATC-3’ site, and triggers the retron-TA, presumably by dissociating the RT-msDNA antitoxin complex (Fig. 2E).

### *racC*-*recE* of Rac prophage are a linked trigger-blocker gene-pair

Exodeoxyribonuclease VIII, encoded by *recE*, a Rac-prophage gene (Fig. 3A), was found to be a retron-TA trigger (Fig. 1B, ED Fig. 1-2). RecE also triggered a retron-TA from an *E. coli* natural isolate (retron-Eco9 ^24^; ED Fig. 6A), suggesting that RecE interacts with a component conserved across the two retrons. RecE is known to specifically recognize double-stranded DNA, and to fully degrade one strand with a 5’→ 3’ directionality ^37^. Since msDNA from all retrons form extended hairpins (dsDNA), we wondered whether the msDNA itself is a RecE-substrate. Indeed, we could not retrieve any msDNA from a strain overexpressing *recE*, while normal levels of msDNA were isolated from a strain carrying the empty-plasmid control (Fig. 3B). This finding suggested that RecE degrades mature msDNA *in vivo*, and thus reduces the functional RT-msDNA antitoxin levels.

Although overexpressing *recE* abolished mature msDNA production, we noticed a higher molecular-weight msDNA-band accumulating (Fig. 3B). This band was of similar molecular weight to that of immature msDNA isolated from Δ*xseAB* strains ^24^, implying that immature msDNA cannot be degraded by RecE *in vivo*. Indeed, overexpressing *recE* in a Δ*xseA* strain yielded similar msDNA levels as when inducing the empty-vector control (Fig. 3C). Nevertheless, this RecE-protection only manifested *in vivo*, since both mature and immature msDNA were cleaved from recombinant RecE *in vitro* (ED Fig. 6B). Since we treated cells with RNase when isolating msDNA, this implies that the RNA-part in the immature msDNA shields the msDNA from RecE.

*recE* is encoded adjacently to a small prophage gene of unknown function, *racC* (Fig. 3A). RacC, a 91 amino acid protein, was the strongest retron-TA blocker identified in the TIC screen (Fig. 1D, ED Fig. 3). RacC blocked RcaT even when expressed without inducer (ED Fig. 4B), and it also blocked the toxin from retron-Eco9 (ED Fig. 6C; albeit requiring higher levels of induction in this case) ^24^. Notably, overexpressing *racC* in *S*Tm completely blocked the RcaT-mediated cold-sensitivity phenotype of all retron antitoxin deletion mutants (Fig. 3E), confirming that RacC acts directly against RcaT activity, rather than through the antitoxin.

In summary, *racC*-*recE* form a linked blocker-trigger gene-pair in the Rac prophage. RecE triggers the retron-TA by directly degrading mature msDNA, and activates RcaT (Fig. 3D), while RacC directly blocks RcaT toxicity, presumably by directly binding to RcaT (Fig. 3F) or by competitive inhibition of the RcaT target.

### A prophage retron-RT triggers the retron-TA by sequestering the msDNA antitoxin

The TransBac overexpression library includes genes from the *E. coli* B strain, which are absent from the *E. coli* K-12 strain ^26^. One of them, RT-Eco1 (B21_00839), is the reverse transcriptase of retron-Eco1 (retron-Ec86), which is encoded in a P2-like prophage ^38^, and was a prominent trigger of the Sen2 retron-TA (Fig. 1C, ED Fig. 1-2). Overexpressing RT-Eco1 in *S*Tm partially activated the endogenous retron-TA, in addition to causing some cold-sensitivity on its own (Fig. 4A, ED Fig. 7). Overexpressing RT-Eco1 also triggered the retron-Eco9 (Fig. 4B), suggesting that RT-Eco1 triggers toxicity by interacting with a conserved retron component.

Since the RT-Eco1 is a retron RT, we wondered whether it disrupts the msDNA biosynthesis of the non-cognate retron-TAs. To test this idea, we isolated msDNA-Sen2 from strains carrying an RT-Eco1 or an empty plasmid. Overexpressing RT-Eco1 did not affect msDNA levels (Fig. 4C). Since the RT-msDNA interaction is essential for antitoxin activity ^24^, we postulated that instead RT-Eco1 could be competing with RT-Sen2, for binding to the mature msDNA-Sen2. This competition would free RT-Sen2 from its cognate msDNA, rendering the antitoxin unit inactive, and therefore activate RcaT. In this case supplying extra copies of msDNA, while maintaining constant protein-levels of the two RTs, would reduce the competition for msDNA-binding, and alleviate the RT-Eco1-mediated triggering of RcaT.

The limiting step in msDNA synthesis are the msrmsd-RNA template-levels, not the RT protein-levels, which produce more msDNA if given more msrmsd-RNA substrate. To supply more msDNA, we overexpressed only the msrmsd-RNA template from a third plasmid. To exclude that RT-Eco1 interacts with the msrmsd-RNA itself, we also supplied msrmsd^mut^ template, which cannot be reverse transcribed into msDNA (due to a mutation in the branching G of the *msr* region ^39^). Indeed, supplementing msrmsd^WT^, but not msrmsd^mut^, completely abolished RT-Eco1 triggering (Fig. 4D). At high IPTG induction-levels, msrmsd^mut^ triggered the retron-TA by itself, presumably because msrmsd^mut^ competes for binding with the native msrmsd template (Fig. 4D). In all cases, toxicity was due to RcaT, since none of the constructs inhibited growth upon co-expressing a p-retron-Δ*rcaT* control vector (Fig. 4D).

Thus, the RT-Eco1 acts as a retron-TA trigger, by competing with the RT-Sen2 and sequestering its non-cognate msDNA-Sen2 from the RT-msDNA antitoxin complex. This activates the toxin RcaT (Fig. 4E).

## DISCUSSION

We have developed a reverse genetics-based method, TIC/TAC, which uses systematic gene overexpression libraries to identify molecular blockers and triggers of TA systems. Overexpression libraries allow us to query the role of genes, which are normally not expressed or kept under tight control in standard laboratory growth conditions, when the TA remains inactive. Such genes are more likely to serve as molecular cues for the TA system. One of the requirements for TIC/TAC is for the studied TA system to work in *E. coli*, or in phylogenetically related enterobacteria, where F-based plasmid conjugation of the overexpression libraries works. The functionality of TA systems from diverse phyla has been routinely assessed in *E. coli* ^13,14,17,40^, hence TIC/TAC can be readily applied to many TA systems. As more overexpression libraries become available, more endogenous molecular cues can be probed. Alternatively, the use of knockdown or knockout libraries ^41–44^ can also provide insights into triggers or blockers, albeit more indirectly. An advantage of using *E. coli* overexpression libraries for TIC/TAC is the unrivaled functional characterization of the *E. coli* genome ^45^, which facilitates mechanistic studies on potential hits.

We applied TIC/TAC to a new tripartite TA encoded by bacterial retron elements: the toxin RcaT is inhibited by protein-protein interactions with an RT-msDNA complex ^24^. We identified multiple triggers and blockers, the majority of which we could validate in targeted assays, confirming that TIC/TAC has few false positives. Some of the identified hits likely point to the target and function of RcaT and/or may feed into the complex biosynthesis of the antitoxin complex ^24^. Among the hits, we noticed that many were phage-related triggers (*dam, recE*, RT-Eco1, B21_00839, *ymfH, tfaP*) and prophage-gene blockers (*racC, dicC, ydaW, yfjH, yjhC*), suggesting an extensive arms-race between the retron-TA and phages. In agreement with our findings, an independent study showed that retrons act as bacterial abortive infection anti-phage defense systems ^46^. We provide insights into the underlying mechanisms through which retron-TAs sense phage attack. In all tested cases, it is the inherent architecture of the tripartite retron-TA systems, and the presence of a DNA component in the antitoxin (msDNA) that enables retrons to sense phages – coupling the sensing to functions needed for phage proliferation.

It is important to note that triggers such as Dam and RecE are also simultaneously anti-restriction genes, used by phages to defend against type II ^47^ or type III ^48^ restriction-modification systems (RM), respectively. Thus, phage defenses against innate bacterial-defense systems (anti-RM proteins), can inadvertently trigger retron-TAs by directly inactivating the RT-msDNA antitoxins, leading to RcaT-mediated abortive infection (Fig. 5). It still remains to be understood why the toxin is active only at cold or anaerobic conditions ^24^ and whether this relates to the specific lifestyle of *S*Tm (e.g., the environment has ambient temperatures and gut is anaerobic), or whether phage infection provides additional cues for RcaT to become active in aerobic conditions at 37°C. Nevertheless, our study suggests the presence of crosstalk between abortive infection systems (Abis) and innate/adaptive anti-phage defenses (RM, CRISPR/Cas), where Abis are directly triggered by phage-products meant to block RM or CRISPR/Cas systems (Fig. 5) ^49^.

How chromosomal-TAs are triggered has been a heavily debated question. The idea that chromosomal-TAs are triggered similarly to plasmid-based TAs, where labile antitoxins are preferentially degraded by stress-induced bacterial proteases, has dominated the field for a long time ^14,50–55^. Yet, chromosomal-TAs do not confer addictive phenotypes to mobile DNA elements ^56–58^, implying that their antitoxins are not labile ^59^. In contrast to non-bound antitoxins, toxin-bound antitoxins are often resistant to protease degradation, and maintain their toxins inactive even under protease-stimulating conditions ^58^. These findings together with the fact that key reports on links between stress, proteases, and TA activation were later discovered to be confounded by biological artefacts ^54,55,60,61^ has reopened the discussion on how chromosomal-TA systems could be activated. Our study provides evidence into how TA systems can be activated by phages. We demonstrate that triggers can directly inactivate the RT-msDNA antitoxin by methylating, degrading, or simply just binding to its msDNA component. This concept is in stark contrast with models where the cue is part of a general stress response (e.g., bacterial proteases or phage-induced host-alterations ^12,62^), which would indirectly induce multiple TA systems. We postulate that more TA systems will have direct and specific cues that interfere with antitoxin or toxin activity, which our TIC/TAC methodology could help to identify. This in turn will propel our understanding of the myriad TA systems found in prokaryotes.

The reasons for multiple TA systems existing in bacterial genomes have also been heavily debated ^63^. In the case of the retron-TA, the trigger-specificity is narrow. For example, Dam-mediated triggering requires that the targeted msDNA contains the 5’-GATC-3’ recognition site in its dsDNA-hairpin sequence. Additionally, msDNA not cleaved by exonuclease VII (immature msDNA) is immune to *recE*-mediated degradation. For most retrons, exonuclease VII is not involved in msDNA biosynthesis ^33^. This finding implies that TA systems (even of the same type) have different triggers, thereby providing one possible explanation for the requirement for high numbers of predicted TAs per genome.

In summary, retron-TAs are tripartite abortive-infection systems. The presence of the hairpin DNA component in the antitoxin enables direct sensing of different facets of essential phage activities. Retrons express thousands of copies of linear ssDNA-hairpins intracellularly (msDNA) ^64^, and encode the antitoxin activity in the RT-msDNA complex ^24^. Applying the TIC/TAC screen developed here to more Abi and TA systems will not only expand our understanding of these fascinating modules, but will also offer new paths for specifically treating bacterial pathogens, either through directly triggering endogenous TAs, or through empowering phage therapy to identify phages that are resistant to such defense systems.

## Supporting information

Supplementary Table 1. Prophage genes enrichment in TIC/TAC hit genes.

Supplementary Table 2. Raw and processed TIC/TAC fitness data.

Supplementary Table 3. Genotypes of bacterial strains used in this study.

Supplementary Table 4. Description of plasmids used in this study.

Supplementary Table 5. Description of construction of plasmids used in this study.

Supplementary Table 6. List of primers used in this study.

## ACKNOWLEDGEMENTS

We thank members of the Typas lab for discussions, and especially Morgane Wartel for assistance in establishing the high-throughput conjugation protocol. This work was supported by the European Molecular Biology Laboratory and the Sofja Kovaleskaja Award of the Alexander von Humboldt Foundation. AM and KM were supported by a fellowship from the EMBL Interdisciplinary Postdoc (EI3POD) programme under Marie Skłodowska-Curie Actions COFUND (grant number 664726). JRE was supported by the National Institutes of Health (K08AI108794). HAP is supported by NIFA (NIFA 2016-11004 & 2017-08881) and DARPA. AT is supported by an ERC consolidator grant, uCARE.

## AUTHOR CONTRIBUTIONS

JRE, MMS, HAP, & AT supervised the study. JB and AT conceived the study. JB, KM, AM & AT designed the experiments. JB, KM & AM performed them. GK analyzed the TIC/TAC screen data. JB, AM & GK designed figures, with inputs from AT. JB & AT wrote the manuscript, with input from all authors.

## DATA AVAILABILITY STATEMENT

All raw/processed data from TIC/TAC screens can be found in Table S2. All unprocessed source images are available upon request.

## CODE AVAILABILITY STATEMENT

The code used for TIC/TAC analysis is available upon request.

## ADDITIONAL INFORMATION

Supplementary information is available for this paper. Correspondence and requests for materials should be addressed to AT.

## METHODS

### Bacterial strains, plasmids, primers, and growth conditions

All bacterial strain genotypes, plasmids, plasmid construction, and primers used in this study are described in Tables S3-S6, respectively. Bacteria were grown in Lysogeny Broth Lennox (LB-Lennox; Tryptone 10 g/L, Yeast Extract 5 g/L, Sodium Chloride 5 g/L). LB-Agar plates (LB plates) were prepared by adding separately-autoclaved 2% molten-Agar in liquid LB. All plasmid-carrying bacterial strains were streaked-out, grown, and assayed with appropriate antibiotics to maintain the plasmids. Plasmids carrying P_BAD_-promoters were induced with 0.2% D-arabinose, while plasmids carrying Ptac promoters were induced with low (0.1 mM) or high (1 mM) concentrations of isopropyl β-D-1-thiogalactopyranoside (IPTG). Bacterial strains with chromosomally-inserted antibiotic-resistance cassettes were streaked-out from stocks on antibiotic-LB plates, but grown/assayed thereafter without antibiotics. Antibiotics used were Spectinomycin (100 μg/mL), Ampicillin (50 μg/mL), Tetracycline (10 μg/mL), and Kanamycin (30 μg/mL). For diaminopimelic acid (DAP) auxotroph-strains, DAP was provided at a final concentration of 0.3 mM. Cold-sensitive strains (*S*Tm retron-Sen2 mutants) were freshly streaked-out from glycerol stocks and kept only at 37°C before every experiment, in order to avoid suppressor mutations. For cold-sensitivity growth tests, strains were incubated at 20°C, or at 15°C, for 48 hours, or 72 hours, respectively.

### Toxin Inhibition/Activation Conjugation (TIC/TAC) procedure

384-colony-arrays of the MOB plasmid library (carried within an *E. coli* F+ strain; JA200 ^25^) and of the TransBac plasmid library (carried within an *E. coli* F^+^ *dapA*^-^ strain; BW38029 ^26^) were pinned from liquid glycerol-stocks to ampicillin-LB and tetracycline-DAP-LB plates, respectively, using a Singer ROTOR and 384-density long-pin Singer RePads, and were grown overnight. Conjugation recipient strains (*E. coli* BW25113 ^66^), carried either a p-*rcaT* plasmid (for TIC), or a p-retron plasmid (for TAC), which both contain a spectinomycin resistance cassette (plasmids detailed in Tables S4-S5). Recipient strains were grown overnight in spectinomycin-LB, and 200 μL of cultures (diluted to OD_595_=0.5) were spread using glass beads on LB plates (for MOB), or on LB-DAP plates (for TransBac). Plates with recipient-lawns were incubated in a non-humid incubator at 37°C for 1 hour. Next, 384-colony-arrays of the donor-libraries were pinned on top of the recipient-lawns, using 384 short-pin Singer RePads. Donor and recipients were allowed to conjugate for 8 hours in a humid incubator at 37°C. Subsequently, cells from the conjugation plates were pinned onto double-antibiotic-selection plates, using 384 short-pin Singer RePads, in order to select for BW25113 transconjugants carrying both plasmids (p-*rcaT*/p-retron + library-plasmids). Double-selection plates contained either ampicillin-spectinomycin, or tetracycline-spectinomycin, for MOB or TransBac libraries, respectively, and transconjugants were grown for 24 hours at 37°C.

Transconjugants were subjected to a second round of selection on double-antibiotic plates, and were also re-arrayed in a 1536-colony format. 1536-colony transconjugant plates were incubated for 10 hours at 37°C, and then each plate was pinned (using 1536-density short-pin Singer RePads) on two replicates of double-antibiotic selection plates (third-round of selection; “source-plates”). Source plates were incubated for 5 hours at 37°C and were used to pin onto double-selection LB-plates (“test-plates”), using 1536-density short-pin RePads. Test-plates contained either no inducer, only arabinose, only IPTG (low or high), or combinations of both (TIC TransBac screen was performed only with low IPTG concentrations). Test-plates were incubated for 13 hours at 37°C, and afterwards imaged using a Canon EOS Rebel T3i camera under controlled light settings (S&P robotics).

### TIC/TAC data analysis

Bacterial colony morphological features for each strain were quantified by using the Iris image-analysis platform ^65^, and colony integral opacity values were used as a fitness proxy. To account the effects of plasmid induction on fitness, we used plates containing only low or high concentrations of IPTG as controls (control plates). These were compared to plates in which the library-plasmids and the p-*rcaT*/p-retron were co-induced with IPTG and arabinose (experiment plates). For quality control, we empirically derived cut-offs for strains that were a) growth-inhibited in the control plates (opacity values < 50,000), b) mucoid in the control plates (colony densities of both replicates > 51 ^65^), and c) noisy strains in control and/or experiment plates (standard deviation for opacity values > 23,000 – median opacities were: TAC control - 103,820, TAC experiment – 71,680, TIC control – 106,941, TIC experiment – 24,357). Strains exceeding any of the three cut-offs were flagged and removed from the final reported dataset, but visible on Table S2. Plate exterior opacity values (four outermost rows and columns) were each multiplicatively corrected to match the mean growth of the interior of the plate. Plate-to-plate biases were also multiplicatively corrected to the same mean. Subsequently, z-scores of those corrected opacity values were calculated per condition, and mean z-scores were calculated per mutant across technical replicates. The final reported score is calculated as the difference between the mean z-scores of each mutant in the experiment and the control plates. All raw and processed data from the TIC/TAC analysis can be found in Table S2.

### TIC/TAC validation procedure

To test candidate genes for blocker/trigger activity, individual conjugation donor strains were single-colony purified from the MOB ^25^ and TransBac ^26^ libraries, and used to construct new transconjugants that were assayed through colony-array and spot growth-tests. To verify that the plasmids contained the appropriate open reading frames, the plasmids were isolated and sequenced.

For colony-array growth tests, MOB donor strains (JA200 ^25^) or TransBac donor strains (BW38029 ^26^) were grown overnight in 600 μL of LB at 37°C in a 1 mL-volume 96-deep-well plate (Thermo Scientific; catalogue number 260251), aliquoted in 96-well plates with 15% glycerol, and kept at -80°C until further use. The 96-colony-array donor strains were conjugated with *E. coli* BW25113 recipient-strains carrying plasmids p-empty, p-retron, p-retron-Δ*rcaT*, and p-*rcaT* in four separate LB agar plates (conjugation as described in the TIC/TAC procedure section). Subsequently, the transconjugant strains were combined in a single 384-array, and pinned on plates containing either no inducer, only arabinose, only IPTG (low or high), or combinations of both. Plates were incubated for 15 hours in a humid-incubator at 37°C, and the 384-density arrayed plates were imaged as described before. Pictures were analysed using Iris ^65^, and integral opacity values were used to calculate fitness scores. Fitness scores for TAC were calculated per condition (per plate) as the opacity ratio between *E. coli* strains carrying p-retron-Δ*rcaT* and p-retron (and trigger-plasmid-X). Fitness scores for TIC were calculated per-IPTG-condition as the opacity ratio between an *E. coli* strain carrying p-*rcaT* (and blocker-plasmid-X) and the average opacity value of strains carrying p-*rcaT* (and trigger-plasmid-X, as negative control).

For spot growth-tests, purified MOB and TransBac library-plasmids were first transformed in MFDpir ^67^ and BW38029 ^26^ conjugation donor-strains, respectively. Next, donors were conjugated with either *E. coli* BW25113 carrying plasmids p-retron-Δ*rcaT* and p-retron (for TAC), or *E. coli* BW25113 carrying plasmid p-*rcaT* (for TIC). Conjugation was carried out as described in the plasmid conjugation section. Strains were spotted on plates containing either no inducer, only arabinose, only IPTG (low or high), or combinations of both, and spot growth tests were carried out as described below.

### Spot growth tests

Single bacterial colonies were inoculated in 2 mL of LB, and incubated at 37°C for 5-6 hours in a roller drum (until an OD_595_ of 5-6). Cultures were then stepwise serially-diluted (ten-fold) eight times in LB (100μL culture + 900μL LB). Subsequently, using a 96-pinner (V&P Scientific, catalogue number: VP 404), ∼10 µL of culture dilutions were spotted on LB plates containing appropriate antibiotics, arabinose, and/or low/high concentrations of IPTG, when applicable. LB plates were grown at appropriate temperatures as described above.

### Genetic techniques

Non-mobilizable plasmids were introduced in *E. coli* BW25113 ^66^ and MFDpir ^67^ strains by TSS transformation ^68^, whereas *S*Tm strains were transformed by electroporation ^69^. Mobilizable plasmids were introduced in bacterial strains by conjugation (described in the Plasmid conjugation section below). To construct the p-retron^mut^ plasmid (*msd*: GATC → GTTC), the source plasmid (p-retron^WT^) was mutagenized through PCR using a kit (NEB; Q5-Site-Directed Mutagenesis Kit, catalogue number E0554S), by following the instructions of the manufacturer. Mutagenic primers used were JB433 and JB434 (primers detailed in Table S6).

### Plasmid conjugation

Mobilizable plasmids were introduced to *E. coli* or *S*Tm strains through conjugation (donors were either *E. coli* JA200 ^25^, *E. coli* BW28029 ^26^ or *E. coli* MFDpir ^67^ strains). Conjugation was carried out by growing single colonies of both recipient and donor strains in LB overnight at 37°C in a drum-roller (LB supplemented with DAP or appropriate antibiotics where applicable). Subsequently, 200 μL of diluted overnight cultures (OD_595_=0.5) of the donor strains were spread on LB-plates (supplemented with DAP if applicable), and plates were incubated at 37°C in a dry incubator for 1 hour. Next, 10 μL of diluted overnight cultures (OD_595_=0.5) of the recipient strains were spotted on top of the lawn of the donor strain, and the conjugation plates were incubated for 6 hours at 37°C. Finally, transconjugant strains were selected by streaking them out from the area where the recipient strains were spotted, in either double-antibiotic selection plates, or single-antibiotic plates but without supplementing them with DAP (if applicable). Selection plates were grown overnight at 37°C, and transconjugants were single-colony purified for further use. The high-throughput conjugation protocol is described in the TIC/TAC procedure section.

### msDNA purification, DpnI/RecE digestion, denaturing electrophoresis, and silver-stain

msDNA was extracted by the alkaline lysis method, as described in ^24^. To assess the methylation status of msDNA, the extracted msDNA was further purified from plasmid DNA contaminants by the crush and soak method ^70^. Briefly, msDNA extracts (from 200 mL of bacterial cultures) were electrophoresed in 12% TBE-polyacrylamide gels (70V for 4.5 hours). The equivalent of extracted msDNA from 60 mL of culture were loaded per well, in order to increase the efficiency of subsequent elution from the acrylamide gel. Gels were stained with ethidium bromide, and the msDNA-containing gel slices were transferred to Eppendorf tubes. Next, the gel-slices were crushed against the walls of the tubes with a tip, suspended in two gel-slice volumes of acrylamide elution buffer ^70^, vortexed, and incubated at 37°C in a table-top roller for 16 hours. Samples were centrifuged at 14,000 rpm/5 min/RT, and the supernatants were transferred to fresh tubes. An equal volume of isopropanol was added to the supernatants, samples were vortexed, and msDNA was precipitated overnight at 4°C. Next, samples were centrifuged at 14,000 rpm/60 min/4°C, pellets were washed once with 1 mL of 70% ethanol, and washed pellets were centrifuged again at 14,000 rpm/60 min/4°C. Pellets (purified msDNA) were air-dried for 15 minutes, and resuspended in 12 μL of distilled water. Subsequently, 1 unit/μL of DpnI (NEB; catalogue number R0176S) was added in 5 μL of purified msDNA and digested overnight at 37°C. In parallel, 5 μL of purified msDNA were incubated in the same buffer (without DpnI), and msDNA-digests were electrophoresed in a denaturing 20% urea-TBE-polyacrylamide gel, as described in ^71^. Briefly, 2x formamide loading buffer (90% formamide, 0.5% EDTA, 0.1% xylene cyanol, 0.1% bromophenol blue) was added to the msDNA-digests, and samples were heated to 95°C for 10 min, and were quickly transferred in ice. The denaturing gel was pre-ran for 1 hour at 55°C, and subsequently msDNA-digests were electrophoresed for 3.5 hours, with constant voltage (60 V). Finally, DNA were stained with silver by using a silver-stain kit (Roth; article number L533.1), following the procedure as described in ^72^.

For digesting msDNA with exodeoxyribonuclease VIII (RecE), msDNA were extracted as described in ^24^, and msDNA extracts were incubated overnight at 37°C with 0.5 units/μL of truncated RecE (NEB; catalogue number M0545S). Subsequently, msDNA-digests were electrophoresed in 12% TBE-Polyacrylamide gels, and stained with ethidium bromide for visualization.

### Statistical analyses

For the analysis of TIC/TAC data, z-score difference means per overexpression strain were calculated as the average z-score difference across clones and replicates, separately for each overexpression library. One-tailed p-values (p.value; Table S2) were calculated for each overexpression strain per induction condition, by using the probability distribution function for the normal distribution. Parameters used were the means and standard deviations of the score distribution of each condition. One-tailed FDR-corrected p-values (q.value; Table S2) were subsequently calculated using the Benjamini-Hochberg method ^73^. For the aggregated prophage genes enrichment analysis (Table S1), hit genes were pooled across the two libraries, and p-values for prophage-gene enrichment were calculated using the Fisher’s exact test. Since both MOB ^25^ and TransBac ^26^ libraries are derived from *E. coli*, only unique genes from the two libraries were accounted for in the background frequency, for the aggregated test. A similar Fisher’s test was also conducted per induction condition, separately for the two libraries (Table S1). Flagged strains were not included in the prophage gene enrichment analyses. *E. coli* BW25113 and *E. coli* BL21 (DE3) genes were annotated as prophage genes by using the PHASTER database ^74^.

## EXTENDED DATA FIGURE LEGENDS

**Extended Data Figure 1.**
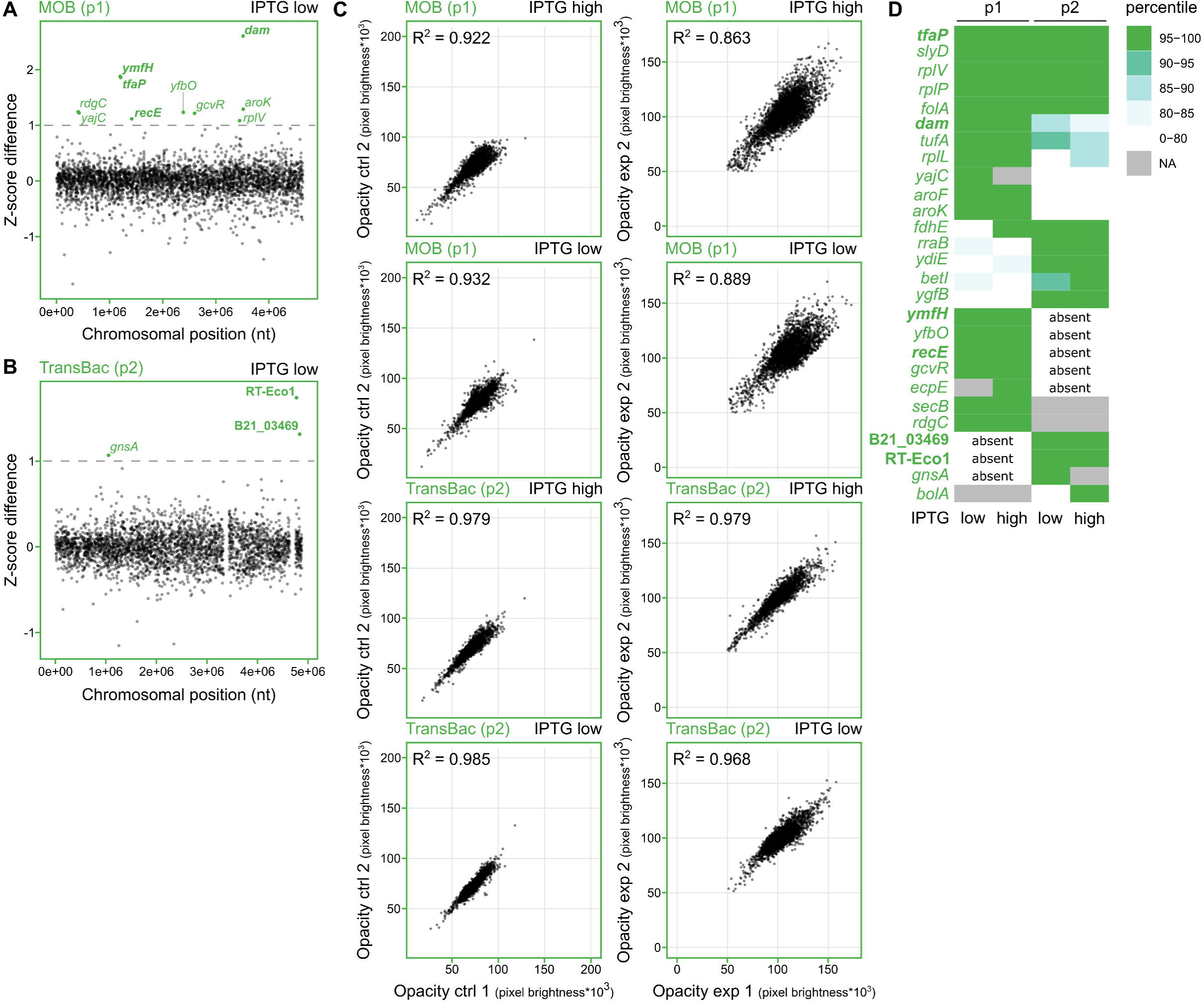
Low-IPTG induction, reproducibility, and hits heatmap for TAC. **(A-B)** TAC results from low-IPTG induction. Experiments as described in Fig. 1B-C, but with low-IPTG plasmid-library induction for MOB (A; p1) ^25^ and TransBac (B; p2) ^26^. **(C)** TAC reproducibility in control and experiment plates. Unprocessed opacity values of strains from control (ctrl) or experiment (exp) plates, derived from replicate plates 1 and 2, were plotted against each other. The overexpression library and IPTG concentration contained in each plate-pair are denoted above the plots. Coefficients of determination (R^2^) are shown for each reproducibility plot. **(D)** Non-parametric comparison of identified triggers across overexpression libraries. Trigger-genes identified from the MOB ^25^ (p1) and the TransBac ^26^ (p2) overexpression libraries were rank-ordered in percentiles based on their mean z-score difference value, as calculated from TAC screens conducted in low, or high-IPTG induction. Not available (NA) denotes genes for which measurements were flagged as problematic (see Methods) in the respective library/IPTG-concentration. Absent denotes genes which are absent from one library. Phage-related trigger-genes are in bold.

**Extended Data Figure 2.**
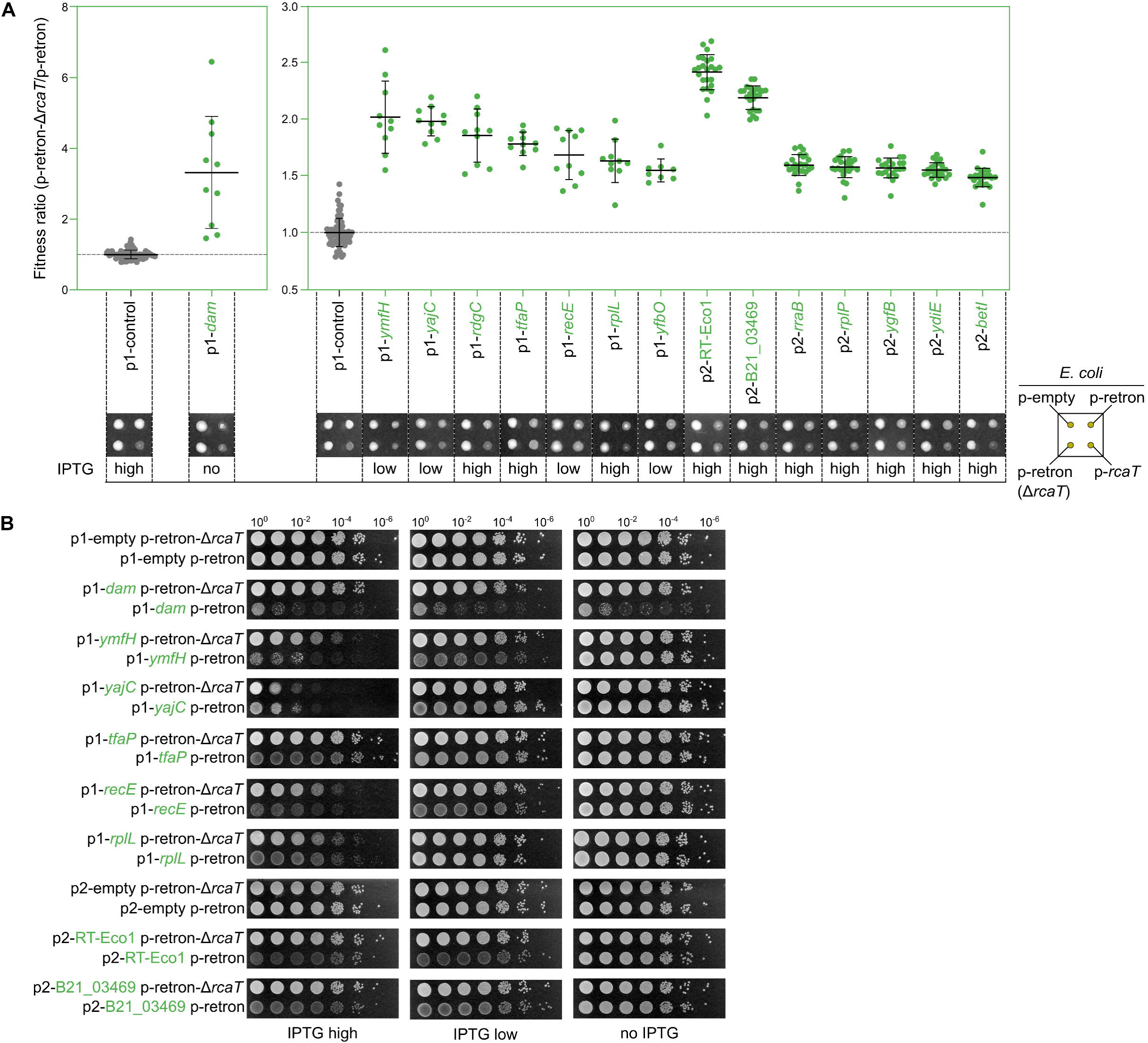
Validation of trigger genes from TAC screens. Triggers specifically inhibit growth by activating RcaT, with triggering degree depending on library-plasmid induction levels. **(A)** Plasmids (p1/p2-trigger gene in green) were conjugated into *E. coli* BW25113 carrying plasmids p-empty, p-retron, p-retron-Δ*rcaT*, or p-*rcaT*. 384-colony-arrays of the transconjugants were pinned on LB plates containing appropriate antibiotics, arabinose, and IPTG concentrations (no IPTG, low, high). The y-axis represents the triggering-degree of each plasmid, measured as the colony opacity ratio of strains (p-retron-Δ*rcaT +* p1/p2-trigger) divided by the (p-retron + p1/p2-trigger). p1-control values were derived by measuring the same colony opacity ratio, for p1-blocker genes (*n*=72; 36 biological X 2 technical replicates), as a negative control. Ratios were calculated from *n*=*10* – 5 biological X 2 technical replicates for p1-trigger genes (except for p1-*yfbO*; *n*=*8* – 4 biological X 2 technical replicates), and from *n=24* – 12 biological X 2 technical replicates for p2-trigger genes. Representative colonies of strains carrying p1/p2-trigger plasmids are shown below the graphs. Horizontal bars denote the average fitness ratio, and error bars denote the standard deviation. Grey horizontal bars denote the p1-control fitness ratio. p1-*dam* was plotted separately, to avoid compressing the scores of the rest of the trigger-genes. **(B)** Plasmids (p1/p2-trigger gene in green, and p1-empty/p2-empty) from conjugation donor strains were conjugated with *E. coli* BW25113 carrying plasmids p-retron-Δ*rcaT* and p-retron. Transconjugants were grown for 5 hours in LB with appropriate antibiotics, serially diluted, spotted on LB plates containing antibiotics, arabinose, and IPTG (no IPTG, low, or high), and incubated at 37°C.

**Extended Data Figure 3.**
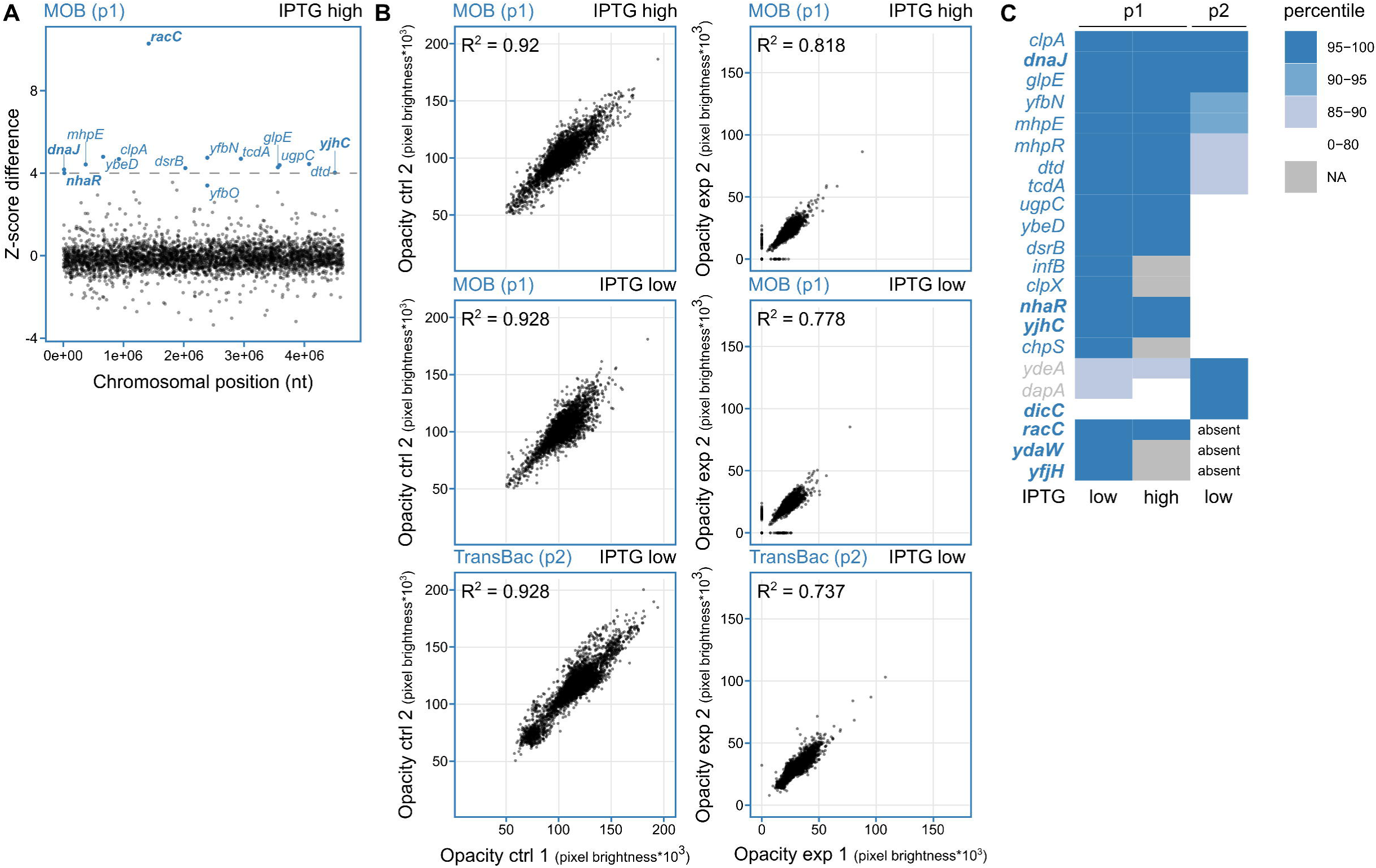
High-IPTG induction, reproducibility, and hits heatmap for TIC. **(A)** TIC results from high-IPTG induction. Experiments as described in Fig. 1D, but with low-IPTG plasmid-library induction for MOB (A; p1) ^25^. Gene *yfbO* was selected as a blocker, despite not passing the significance cut-off, due to genetic linkage with *yfbN*. **(B)** TIC reproducibility in control and experiment plates – as in EDFig. 1C. **(C)** Non-parametric comparison of identified blockers across overexpression libraries – as in EDFig. 1D. Grey-colored genes are presumably affecting induction and/or conjugation levels. Phage-related trigger-genes are in bold.

**Extended Data Figure 4.**
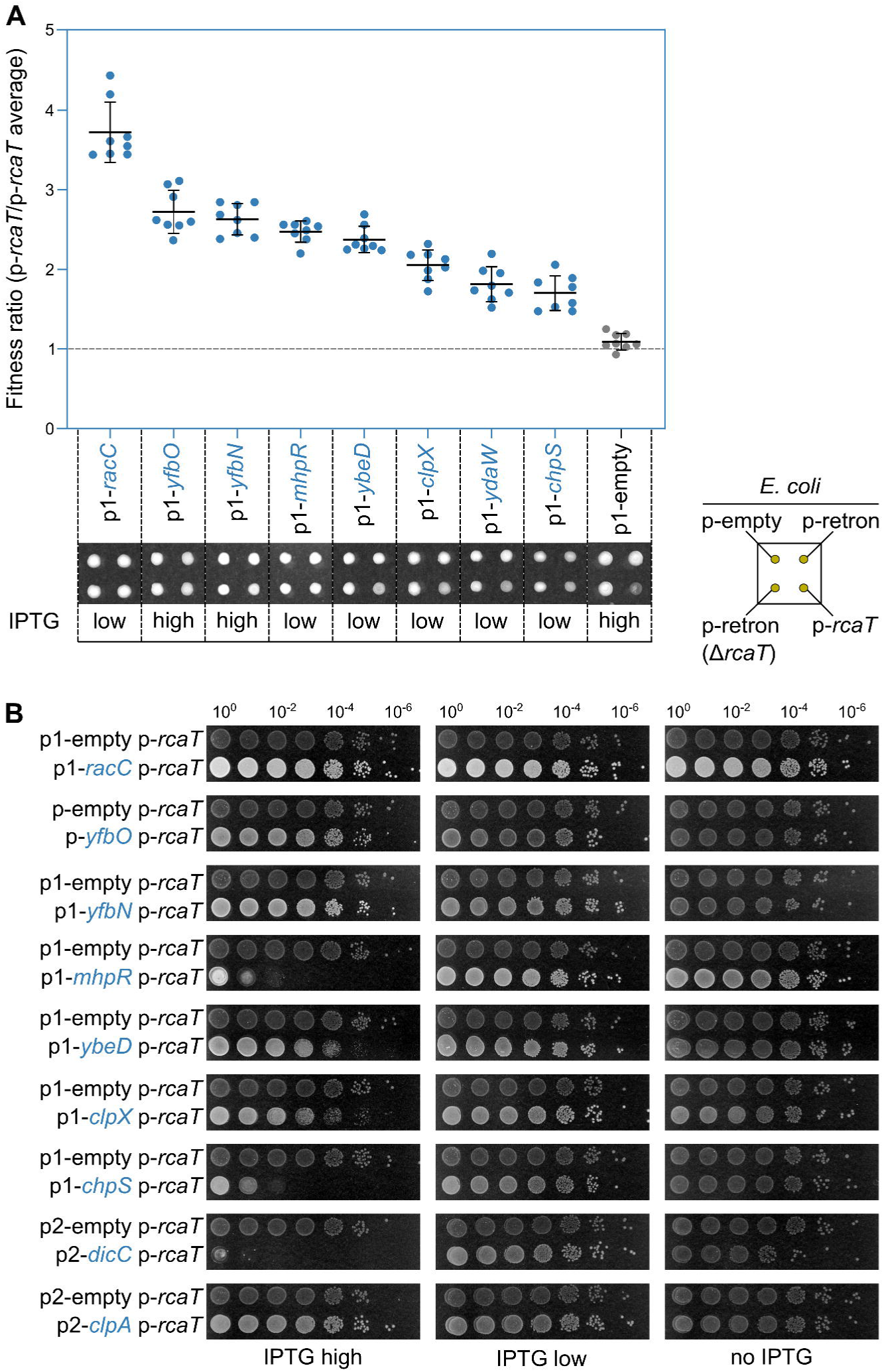
Validation of blocker genes from TIC screens. Blockers specifically alleviate RcaT-mediated toxicity with blocking degree depending on library-plasmid induction levels. **(A)** Plasmids (p1-blocker gene in blue and p1-empty in black) were conjugated into *E. coli* BW25113 carrying plasmids p-empty, p-retron, p-retron-Δ*rcaT*, or p-*rcaT*. 384-colony-arrays of the transconjugants were pinned on LB plates containing appropriate antibiotics, arabinose, and IPTG concentrations (no IPTG, low, or high). The y-axis represents the blocking degree of each plasmid, measured as the colony opacity ratio of strains carrying the different blockers (p-*rcaT +* p1-blocker) divided by the average of a strain carrying trigger plasmids (p-*rcaT* + p1-trigger), as a negative control. Fitness ratios calculated from *n=8* – 4 biological X 2 technical replicates. The mean colony opacity for p1-trigger was derived from *n*=110 – 55 biological X 2 technical replicates. Representative colonies of strains carrying p1-blocker plasmids shown below the graphs. Horizontal bars denote the average fitness ratio, and error bars denote the standard deviation. Grey horizontal bar denotes the expected fitness ratio in the absence of blocking effects. **(B)** Plasmids (p1/p2-blocker gene in blue, and p1-empty/p2-empty) from conjugation donor strains were conjugated with *E. coli* BW25113 carrying plasmid p-*rcaT* or a p1-empty plasmid. Transconjugants were grown as in EDFig. 2B.

**Extended Data Figure 5.**
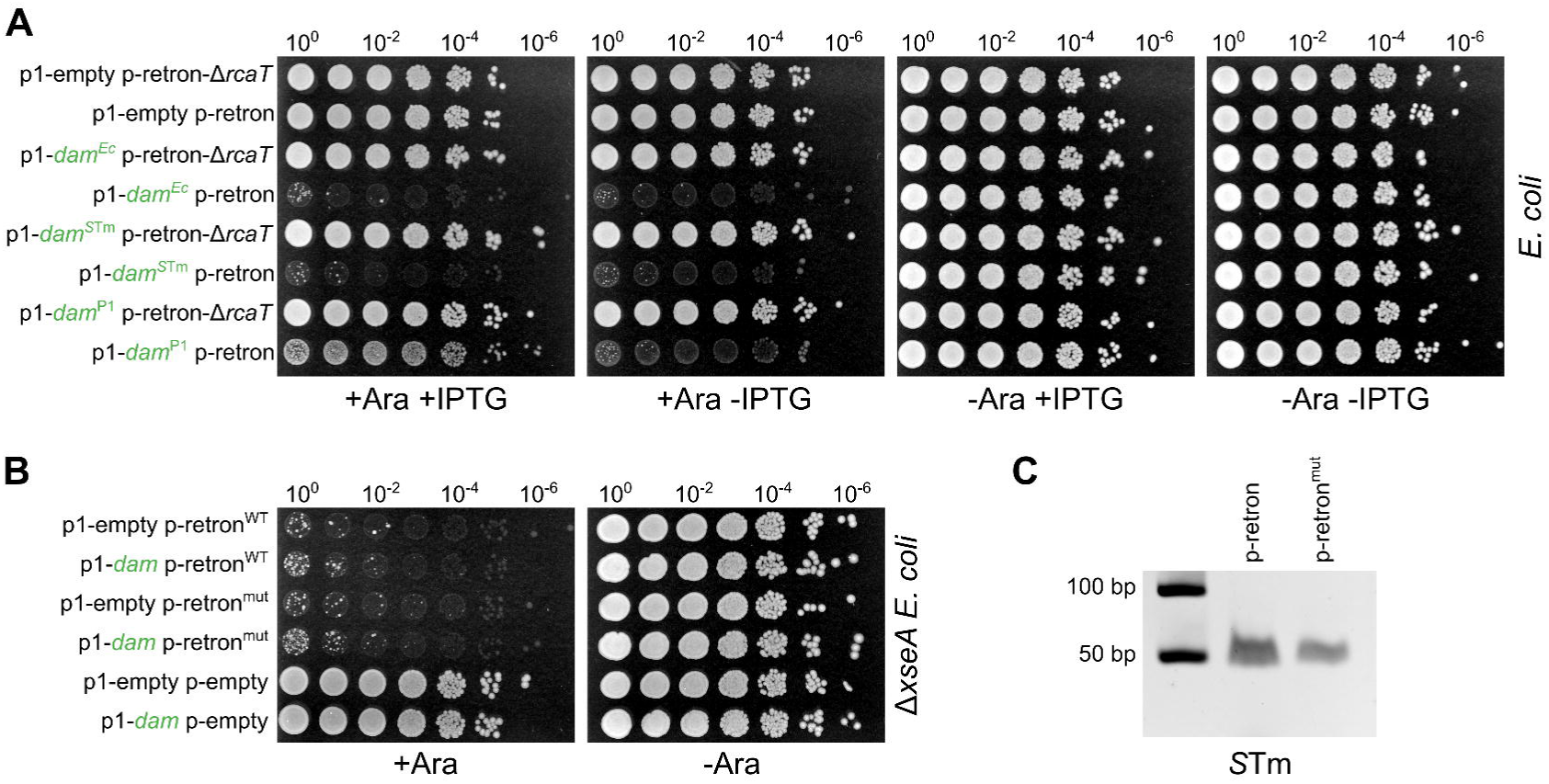
Dam-mediated retron-TA triggering. **(A)** Bacterial/phage *dam* homologues trigger the retron-TA. *E. coli* BW25113 carrying combinations of plasmids p-retron-Δ*rcaT*, p-retron, p1-*dam*^Ec^, p1-*dam*^*S*Tm^, p1-*dam*^P1^, and p1-empty, were grown for 5 hours in LB with appropriate antibiotics, serially diluted, spotted on LB plates with antibiotics, with/without arabinose, and IPTG (low), and incubated at 37°C. **(B)** *rcaT* in p-retron^mut^ is functional and expressed. *E. coli* BW25113 Δ*xseA* strains carrying combinations of plasmids p-retron^WT^, p-retron^mut^, p1-*dam*, and p1-empty, were grown and spotted as described in panel **A**. Representative data shown from two independent experiments. **(C)** p-retron^mut^ produces msDNA at equivalent levels as wildtype retron. msDNA was extracted from *S*Tm strains carrying p-retron^WT^, or p-retron^mut^. Plasmids were induced with arabinose. Extracted msDNA was electrophoresed in a TBE-Polyacrylamide gel.

**Extended Data Figure 6.**
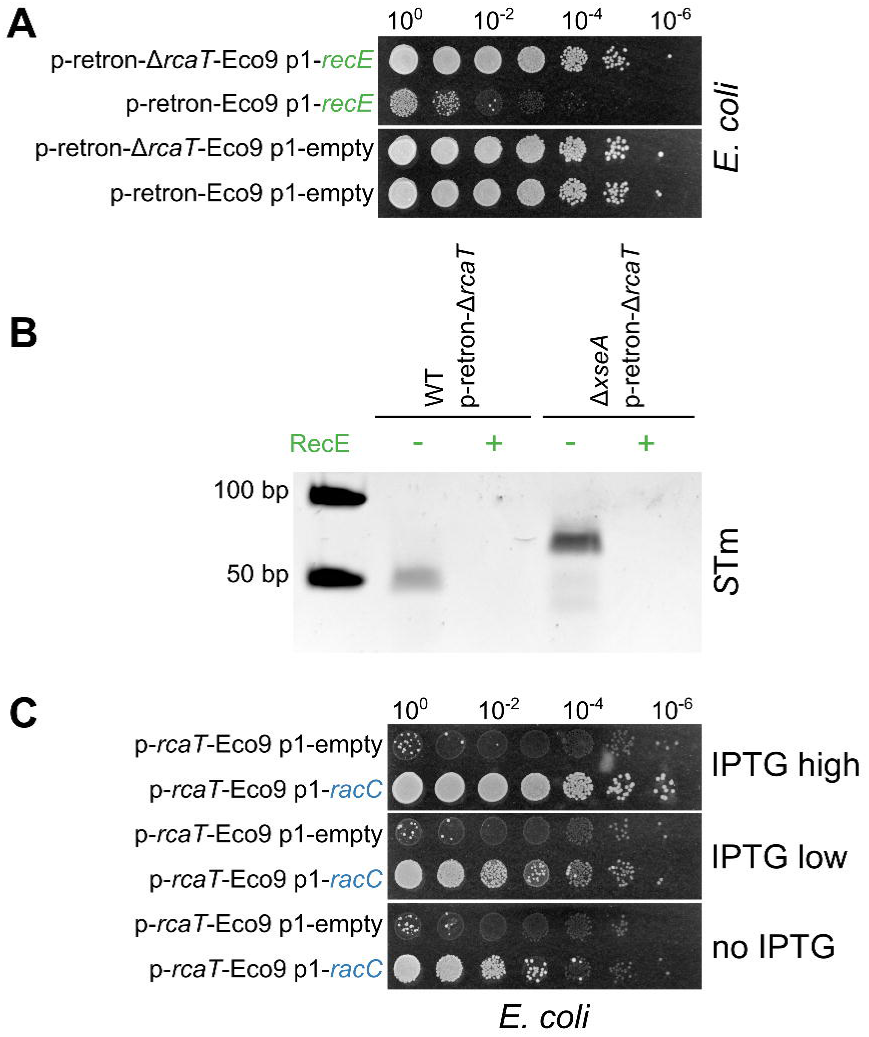
*racC*-*recE* constitute a blocker-trigger gene-pair for retron-Eco9. **(A)** RecE triggers the retron-Eco9. *E. coli* BW25113 carrying combinations of plasmids p-retron-Eco9, p-retron-Δ*rcaT*-Eco9, p1-*recE*, and p1-empty, were grown for 5 hours in LB with appropriate antibiotics, serially diluted, spotted on LB plates with antibiotics, arabinose, and IPTG (low), and incubated at 37°C. Representative data shown from three independent experiments. **(B)** RecE degrades mature and immature msDNA *in vitro*. msDNA was extracted from *S*Tm strains (WT, or Δ*xseA*) carrying plasmid p-retron-Δ*rcaT*. msDNA extracts were incubated with recombinant RecE (see Methods), and digests were electrophoresed on TBE-Polyacrylamide gels. **(C)** RacC also blocks RcaT-Eco9. *E. coli* BW25113 carrying combinations of plasmids p-*rcaT*-Eco9, p1-*racC*, and p1-empty, were grown and spotted as in panel **A** with different IPTG concentrations.

**Extended Data Figure 7.**
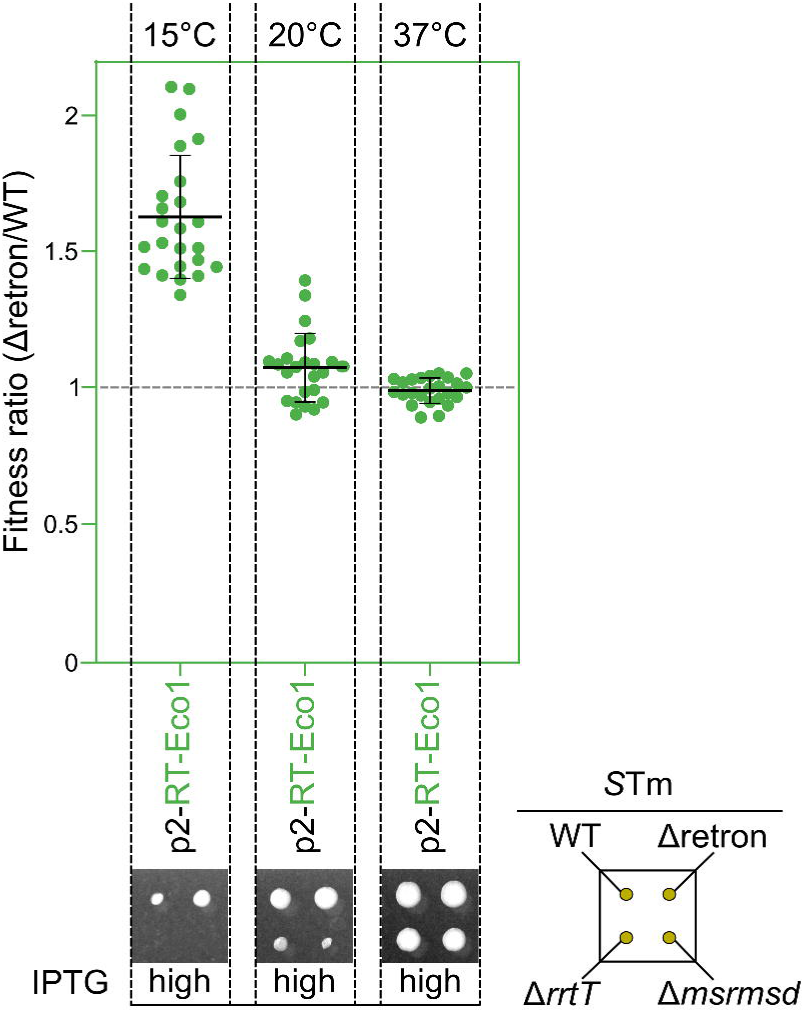
RT-Eco1 partially triggers endogenous *S*Tm retron-TA. *S*Tm strains (WT, Δretron, Δ*rrtT*, Δ*msrmsd*) carrying plasmid p2-RT-Eco1 were arrayed on LB plates containing tetracycline and IPTG (high), and plates were incubated either at 15°C, 20°C, or 37°C. The y-axis represents the triggering degree of RT-Eco1, measured as the colony size ratio of strain (Δretron *+* p2-RT-Eco1) divided by the (WT + p2-RT-Eco1). Fitness ratios calculated from *n*=24 – 12 biological ł 2 technical replicates. Representative *S*Tm colonies carrying p2-RT-Eco1 are shown below the graph. Horizontal bars denote the average fitness ratio, and error bars denote standard deviation. Grey bar denotes the expected fitness ratio in the absence of triggering effects.

## SUPPLEMENTARY INFORMATION

**Supplementary Table 1**. Prophage genes enrichment in TIC/TAC hit genes.

**Supplementary Table 2**. Raw and processed TIC/TAC fitness data.

**Supplementary Table 3**. Genotypes of bacterial strains used in this study.

**Supplementary Table 4**. Description of plasmids used in this study.

**Supplementary Table 5**. Description of construction of plasmids used in this study.

**Supplementary Table 6**. List of primers used in this study.

