## Supplementary Table 3. Genotypes of bacterial strains used in this study. for "Phage proteins block and trigger retron toxin/antitoxin systems"

**Supplementary Table 3.** Genotypes of bacterial strains used in this study (1/3).

| **Name (as in the figures)** | **Genotype** | **Strain code** | **Used in** | **Used for** |
|---|---|---|---|---|
| *S*Tm | *Salmonella enterica* enterica Typhimurium 14028s | *S*Tm | - | Wildtype *S*Tm |
| *E. coli* | *Escherichia coli* BW25113 | *E. coli* | - | Wildtype *E. coli* |
| p1-empty | *S*Tm Δ*araBAD*::kan pNTR-SD | NT16721 | Fig. 2A | Cold-sensitivity growth assay |
| p1-*dam* | *S*Tm Δ*araBAD*::kan pNTR-SD-*dam* | NT16820 | Fig. 2A | Cold-sensitivity growth assay |
| p1-empty Δ*rcaT* | *S*Tm ΔrcaT::FRT ΔaraBAD::kan pNTR-SD | NT16993 | Fig. 2A | Cold-sensitivity growth assay |
| p1-*dam* Δ*rcaT* | *S*Tm ΔrcaT::FRT Δ*araBAD*::kan pNTR-SD | NT16727 | Fig. 2A | Cold-sensitivity growth assay |
| p-retron-Δ*rcaT* p1-empty | *S*Tm Δretron::FRT Δ*araBAD*::kan pNTR-SD | NT32175 | Fig. 2B | msDNA isolation |
| p-retron-Δ*rcaT* p1-*dam* | *S*Tm Δretron::FRT Δ*araBAD*::kan pNTR-SD-*dam* | NT32176 | Fig. 2B | msDNA isolation |
| p1-empty p-retron$^{WT}$ | *E. coli* pNTR-SD pJB38 | NT32167 | Fig. 2C | Retron over-expression growth assay |
| p1-*dam* p-retron$^{WT}$ | *E. coli* pNTR-SD-*dam* pJB38 | NT32169 | Fig. 2C | Retron over-expression growth assay |
| p1-empty p-retron$^{mut}$ | *E. coli* pNTR-SD pJB128 | NT32171 | Fig. 2C | Retron over-expression growth assay |
| p1-*dam* p-retron$^{mut}$ | *E. coli* pNTR-SD-*dam* pJB128 | NT32173 | Fig. 2C | Retron over-expression growth assay |
| p1-empty p-empty | *E. coli* pNTR-SD pTU175 | NT32163 | Fig. 2C | Retron over-expression growth assay |
| p1-*dam* p-empty | *E. coli* pNTR-SD-*dam* pTU175 | NT32165 | Fig. 2C | Retron over-expression growth assay |
| msDNA (WT) p1-*dam* (-) | *S*Tm Δretron::FRT Δ*araBAD*::kan pNTR-SD pJB51 | NT32179 | Fig. 2D | msDNA purification to cleave with DpnI |
| msDNA (WT) p1-*dam* (+) | *S*Tm Δretron::FRT Δ*araBAD*::kan pNTR-SD-*dam* pJB51 | NT32181 | Fig. 2D | msDNA purification to cleave with DpnI |
| msDNA (mut) p1-*dam* (-) | *S*Tm Δretron::FRT Δ*araBAD*::kan pNTR-SD pJB128 | NT32180 | Fig. 2D | msDNA purification to cleave with DpnI |
| msDNA (mut) p1-*dam* (+) | *S*Tm Δretron::FRT Δ*araBAD*::kan pNTR-SD-*dam* pJB128 | NT32182 | Fig. 2D | msDNA purification to cleave with DpnI |
| WT p-retron-Δ*rcaT* p1-empty | *S*Tm Δretron::FRT Δ*araBAD*::kan pNTR-SD | NT32175 | Fig. 3B | msDNA isolation |
| WT p-retron-Δ*rcaT* p1-*recE* | *S*Tm Δretron::FRT Δ*araBAD*::kan pNTR-SD-*recE* | NT16684 | Fig. 3B | msDNA isolation |
| Δ*xseA* p-retron-Δ*rcaT* p1-empty | *S*Tm Δ*xseA*::FRT Δ*araBAD*::kan pNTR-SD | NT16723 | Fig. 3C | msDNA isolation |
| Δ*xseA* p-retron-Δ*rcaT* p1-*recE* | *S*Tm Δ*xseA*::FRT Δ*araBAD*::kan pNTR-SD-*recE* | NT16686 | Fig. 3C | msDNA isolation |
| p1-empty | *S*Tm Δ*araBAD*::kan pNTR-SD | NT16721 | Fig. 3E | Cold-sensitivity growth assay |
| p1-*racC* | *S*Tm Δ*araBAD*::kan pNTR-SD-*racC* | NT16827 | Fig. 3E | Cold-sensitivity growth assay |
| Δ*rrtT* p1-empty | *S*Tm Δ*rrtT*::FRT Δ*araBAD*::kan pNTR-SD | NT16722 | Fig. 3E | Cold-sensitivity growth assay |
| Δ*rrtT* p1-*racC* | *S*Tm Δ*rrtT*::FRT Δ*araBAD*::kan pNTR-SD-*racC* | NT16828 | Fig. 3E | Cold-sensitivity growth assay |
| Δ*xseA* p1-empty | *S*Tm Δ*xseA*::FRT Δ*araBAD*::kan pNTR-SD | NT16723 | Fig. 3E | Cold-sensitivity growth assay |
| Δ*xseA* p1-*racC* | *S*Tm Δ*xseA*::FRT Δ*araBAD*::kan pNTR-SD-*racC* | NT16829 | Fig. 3E | Cold-sensitivity growth assay |
| Δ*rnhA* p1-empty | *S*Tm Δ*rnhA*::FRT Δ*araBAD*::kan pNTR-SD | NT16725 | Fig. 3E | Cold-sensitivity growth assay |
| Δ*rnhA* p1-*racC* | *S*Tm Δ*rnhA*::FRT Δ*araBAD*::kan pNTR-SD-*racC* | NT16831 | Fig. 3E | Cold-sensitivity growth assay |
| Δ*msrmsd* p1-empty | *S*Tm Δ*msrmsd*::FRT Δ*araBAD*::kan pNTR-SD | NT16726 | Fig. 3E | Cold-sensitivity growth assay |
| Δ*msrmsd* p1-*racC* | *S*Tm Δ*msrmsd*::FRT Δ*araBAD*::kan pNTR-SD-*racC* | NT16833 | Fig. 3E | Cold-sensitivity growth assay |
| p2-empty | *S*Tm Δ*araBAD*::kan pFE604T | NT16876 | Fig. 4A | Cold-sensitivity growth assay |
| p2-RT-Eco1 | *S*Tm Δ*araBAD*::kan pFE604T-B21_00839 | NT16834 | Fig. 4A | Cold-sensitivity growth assay |
| Δ*rcaT* p2-empty | *S*Tm Δ*rcaT*::FRT Δ*araBAD*::kan pFE604T | NT16880 | Fig. 4A | Cold-sensitivity growth assay |
| Δ*rcaT* p2-RT-Eco1 | *S*Tm Δ*rcaT*::FRT Δ*araBAD*::kan pFE604T-B21_00839 | NT16839 | Fig. 4A | Cold-sensitivity growth assay |
| p-retron-ΔrcaT-Eco9 p2-RT-Eco1 | *E. coli* pJB95 pFE604T-B21_00839 | - | Fig. 4B | RcaT-Eco9 over-expression growth assay |
| p-retron-Eco9 p2-RT-Eco1 | *E. coli* pJB118 pFE604T-B21_00839 | - | Fig. 4B | RcaT-Eco9 over-expression growth assay |
| p-retron-ΔrcaT-Eco9 p2-empty | *E. coli* pJB95 pFE604T | - | Fig. 4B | RcaT-Eco9 over-expression growth assay |
| p-retron-Eco9 p2-empty | *E. coli* pJB118 pFE604T | - | Fig. 4B | RcaT-Eco9 over-expression growth assay |
| p-retron-Δ*rcaT* p2-empty | *S*Tm Δretron::FRT Δ*araBAD*::kan pJB51 pFE604T | NT16865 | Fig. 4C | msDNA isolation |
| p-retron-Δ*rcaT* p2-RT-Eco1 | *S*Tm Δretron::FRT Δ*araBAD*::kan pJB51 pFE604T-B21_00839 | NT16864 | Fig. 4C | msDNA isolation |
| p-retron p1-msrmsd (-) p2-RT-Eco1 (-) | *E. coli* pJB38 pNTR-SD pFE604T | NT32217 | Fig. 4D | Retron over-expression growth assay |
| p-retron p1-msrmsd (-) p2-RT-Eco1 (+) | *E. coli* pJB38 pNTR-SD pFE604T-B21_00839 | NT32220 | Fig. 4D | Retron over-expression growth assay |
| p-retron p1-msrmsd (WT) p2-RT-Eco1 (-) | *E. coli* pJB38 pKM1 pFE604T | NT32218 | Fig. 4D | Retron over-expression growth assay |
| p-retron p1-msrmsd (WT) p2-RT-Eco1 (+) | *E. coli* pJB38 pKM1 pFE604T-B21_00839 | NT32221 | Fig. 4D | Retron over-expression growth assay |
| p-retron p1-msrmsd (mut) p2-RT-Eco1 (-) | *E. coli* pJB38 pKM3 pFE604T | NT32219 | Fig. 4D | Retron over-expression growth assay |
| p-retron p1-msrmsd (mut) p2-RT-Eco1 (+) | *E. coli* pJB38 pKM3 pFE604T-B21_00839 | NT32222 | Fig. 4D | Retron over-expression growth assay |
| p-retron-Δ*rcaT* p1-msrmsd (-) p2-RT-Eco1 (-) | *E. coli* pJB51 pNTR-SD pFE604T | NT32211 | Fig. 4D | Retron over-expression growth assay |
| p-retron-Δ*rcaT* p1-msrmsd (-) p2-RT-Eco1 (+) | *E. coli* pJB51 pNTR-SD pFE604T-B21_00839 | NT32214 | Fig. 4D | Retron over-expression growth assay |
| p-retron-Δ*rcaT* p1-msrmsd (WT) p2-RT-Eco1 (-) | *E. coli* pJB51 pKM1 pFE604T | NT32212 | Fig. 4D | Retron over-expression growth assay |
| p-retron-Δ*rcaT* p1-msrmsd (WT) p2-RT-Eco1 (+) | *E. coli* pJB51 pKM1 pFE604T-B21_00839 | NT32215 | Fig. 4D | Retron over-expression growth assay |
| p-retron-Δ*rcaT* p1-msrmsd (mut) p2-RT-Eco1 (-) | *E. coli* pJB51 pKM3 pFE604T | NT32213 | Fig. 4D | Retron over-expression growth assay |
| p-retron-Δ*rcaT* p1-msrmsd (mut) p2-RT-Eco1 (+) | *E. coli* pJB51 pKM3 pFE604T-B21_00839 | NT32216 | Fig. 4D | Retron over-expression growth assay |
| p1-*dam* p-empty | *E. coli* pTU175 pNTR-SD-*dam* | - | ED Fig. 2A | Retron over-expression growth assay (trigger validation) |
| p1-*dam* p-retron | *E. coli* pJB38 pNTR-SD-*dam* | - | ED Fig. 2A/2B | Retron over-expression growth assay (trigger validation) |
| p1-*dam* p-retron-Δ*rcaT* | *E. coli* pJB51 pNTR-SD-*dam* | - | ED Fig. 2A/2B | Retron over-expression growth assay (trigger validation) |
| p1-*dam* p-*rcaT* | *E. coli* pJB37 pNTR-SD-*dam* | - | ED Fig. 2A | Retron over-expression growth assay (trigger validation) |
| p1-*ymfH* p-empty | *E. coli* pTU175 pNTR-SD-*ymfH* | - | ED Fig. 2A | Retron over-expression growth assay (trigger validation) |
| p1-*ymfH* p-retron | *E. coli* pJB38 pNTR-SD-*ymfH* | - | ED Fig. 2A/2B | Retron over-expression growth assay (trigger validation) |
| p1-*ymfH* p-retron-Δ*rcaT* | *E. coli* pJB51 pNTR-SD-*ymfH* | - | ED Fig. 2A/2B | Retron over-expression growth assay (trigger validation) |
| p1-*ymfH* p-*rcaT* | *E. coli* pJB37 pNTR-SD-*ymfH* | - | ED Fig. 2A | Retron over-expression growth assay (trigger validation) |
| p1-*yajC* p-empty | *E. coli* pTU175 pNT3-*yajC* | - | ED Fig. 2A | Retron over-expression growth assay (trigger validation) |
| p1-*yajC* p-retron | *E. coli* pJB38 pNT3-*yajC* | - | ED Fig. 2A/2B | Retron over-expression growth assay (trigger validation) |
| p1-*yajC* p-retron-Δ*rcaT* | *E. coli* pJB51 pNT3-*yajC* | - | ED Fig. 2A/2B | Retron over-expression growth assay (trigger validation) |

**Supplementary Table 3.** Genotypes of bacterial strains used in this study (2/3).

| | | | | |
|---|---|---|---|---|
| p1-*yajC* p-*rcaT* | *E. coli* pJB37 pNT3-*yajC* | - | ED Fig. 2A | Retron over-expression growth assay (trigger validation) |
| p1-*rdgC* p-empty | *E. coli* pTU175 pNTR-SD-*rdgC* | - | ED Fig. 2A | Retron over-expression growth assay (trigger validation) |
| p1-*rdgC* p-retron | *E. coli* pJB38 pNTR-SD-*rdgC* | - | ED Fig. 2A | Retron over-expression growth assay (trigger validation) |
| p1-*rdgC* p-retron-Δ*rcaT* | *E. coli* pJB51 pNTR-SD-*rdgC* | - | ED Fig. 2A | Retron over-expression growth assay (trigger validation) |
| p1-*rdgC* p-*rcaT* | *E. coli* pJB37 pNTR-SD-*rdgC* | - | ED Fig. 2A | Retron over-expression growth assay (trigger validation) |
| p1-*tfaP* p-empty | *E. coli* pTU175 pNT3-*tfaP* | - | ED Fig. 2A | Retron over-expression growth assay (trigger validation) |
| p1-*tfaP* p-retron | *E. coli* pJB38 pNT3-*tfaP* | - | ED Fig. 2A/2B | Retron over-expression growth assay (trigger validation) |
| p1-*tfaP* p-retron-Δ*rcaT* | *E. coli* pJB51 pNT3-*tfaP* | - | ED Fig. 2A/2B | Retron over-expression growth assay (trigger validation) |
| p1-*tfaP* p-*rcaT* | *E. coli* pJB37 pNT3-*tfaP* | - | ED Fig. 2A | Retron over-expression growth assay (trigger validation) |
| p1-*recE* p-empty | *E. coli* pTU175 pNTR-SD-*recE* | - | ED Fig. 2A | Retron over-expression growth assay (trigger validation) |
| p1-*recE* p-retron | *E. coli* pJB38 pNTR-SD-*recE* | - | ED Fig. 2A/2B | Retron over-expression growth assay (trigger validation) |
| p1-*recE* p-retron-Δ*rcaT* | *E. coli* pJB51 pNTR-SD-*recE* | - | ED Fig. 2A/2B | Retron over-expression growth assay (trigger validation) |
| p1-*recE* p-*rcaT* | *E. coli* pJB37 pNTR-SD-*recE* | - | ED Fig. 2A | Retron over-expression growth assay (trigger validation) |
| p1-*ygaV* p-empty | *E. coli* pTU175 pNT3-*ygaV* | - | ED Fig. 2A | Retron over-expression growth assay (trigger validation) |
| p1-*ygaV* p-retron | *E. coli* pJB38 pNT3-*ygaV* | - | ED Fig. 2A | Retron over-expression growth assay (trigger validation) |
| p1-*ygaV* p-retron-Δ*rcaT* | *E. coli* pJB51 pNT3-*ygaV* | - | ED Fig. 2A | Retron over-expression growth assay (trigger validation) |
| p1-*ygaV* p-*rcaT* | *E. coli* pJB37 pNT3-*ygaV* | - | ED Fig. 2A | Retron over-expression growth assay (trigger validation) |
| p1-*rplL* p-empty | *E. coli* pTU175 pNT3-*rplL* | - | ED Fig. 2A | Retron over-expression growth assay (trigger validation) |
| p1-*rplL* p-retron | *E. coli* pJB38 pNT3-*rplL* | - | ED Fig. 2A/2B | Retron over-expression growth assay (trigger validation) |
| p1-*rplL* p-retron-Δ*rcaT* | *E. coli* pJB51 pNT3-*rplL* | - | ED Fig. 2A/2B | Retron over-expression growth assay (trigger validation) |
| p1-*rplL* p-*rcaT* | *E. coli* pJB37 pNT3-*rplL* | - | ED Fig. 2A | Retron over-expression growth assay (trigger validation) |
| p1-*yfbO* p-empty | *E. coli* pTU175 pNT3-*yfbO* | - | ED Fig. 2A | Retron over-expression growth assay (trigger validation) |
| p1-*yfbO* p-retron | *E. coli* pJB38 pNT3-*yfbO* | - | ED Fig. 2A | Retron over-expression growth assay (trigger validation) |
| p1-*yfbO* p-retron-Δ*rcaT* | *E. coli* pJB51 pNT3*yfbO* | - | ED Fig. 2A | Retron over-expression growth assay (trigger validation) |
| p1-*yfbO* p-*rcaT* | *E. coli* pJB37 pNT3-*yfbO* | - | ED Fig. 2A | Retron over-expression growth assay (trigger validation) |
| p2-RT-Eco1 p-empty | *E. coli* pTU175 pFE604T-B21_00839 | - | ED Fig. 2A | Retron over-expression growth assay (trigger validation) |
| p2-RT-Eco1 p-retron | *E. coli* pJB38 pFE604T-B21_00839 | - | ED Fig. 2A/2B | Retron over-expression growth assay (trigger validation) |
| p2-RT-Eco1 p-retron-Δ*rcaT* | *E. coli* pJB51 pFE604T-B21_00839 | - | ED Fig. 2A/2B | Retron over-expression growth assay (trigger validation) |
| p2-RT-Eco1 p-*rcaT* | *E. coli* pJB37 pFE604T-B21_00839 | - | ED Fig. 2A | Retron over-expression growth assay (trigger validation) |
| p2-B21_03469 p-empty | *E. coli* pTU175 pFE604T-B21_03469 | - | ED Fig. 2A | Retron over-expression growth assay (trigger validation) |
| p2-B21_03469 p-retron | *E. coli* pJB38 pFE604T-B21_03469 | - | ED Fig. 2A/2B | Retron over-expression growth assay (trigger validation) |
| p2-B21_03469 p-retron-Δ*rcaT* | *E. coli* pJB51 pFE604T-B21_03469 | - | ED Fig. 2A/2B | Retron over-expression growth assay (trigger validation) |
| p2-B21_03469 p-*rcaT* | *E. coli* pJB37 pFE604T-B21_03469 | - | ED Fig. 2A | Retron over-expression growth assay (trigger validation) |
| p2-*rraB* p-empty | *E. coli* pTU175 pFE604T-*rraB* | - | ED Fig. 2A | Retron over-expression growth assay (trigger validation) |
| p2-*rraB* p-retron | *E. coli* pJB38 pFE604T-*rraB* | - | ED Fig. 2A | Retron over-expression growth assay (trigger validation) |
| p2-*rraB* p-retron-Δ*rcaT* | *E. coli* pJB51 pFE604T-*rraB* | - | ED Fig. 2A | Retron over-expression growth assay (trigger validation) |
| p2-*rraB* p-*rcaT* | *E. coli* pJB37 pFE604T-*rraB* | - | ED Fig. 2A | Retron over-expression growth assay (trigger validation) |
| p2-*ygfB* p-empty | *E. coli* pTU175 pFE604T-*ygfB* | - | ED Fig. 2A | Retron over-expression growth assay (trigger validation) |
| p2-*ygfB* p-retron | *E. coli* pJB38 pFE604T-*ygfB* | - | ED Fig. 2A | Retron over-expression growth assay (trigger validation) |
| p2-*ygfB* p-retron-Δ*rcaT* | *E. coli* pJB51 pFE604T-*ygfB* | - | ED Fig. 2A | Retron over-expression growth assay (trigger validation) |
| p2-*ygfB* p-*rcaT* | *E. coli* pJB37 pFE604T-*ygfB* | - | ED Fig. 2A | Retron over-expression growth assay (trigger validation) |
| p2-*ydiE* p-empty | *E. coli* pTU175 pFE604T-*ydiE* | - | ED Fig. 2A | Retron over-expression growth assay (trigger validation) |
| p2-*ydiE* p-retron | *E. coli* pJB38 pFE604T-*ydiE* | - | ED Fig. 2A | Retron over-expression growth assay (trigger validation) |
| p2-*ydiE* p-retron-Δ*rcaT* | *E. coli* pJB51 pFE604T-*ydiE* | - | ED Fig. 2A | Retron over-expression growth assay (trigger validation) |
| p2-*ydiE* p-*rcaT* | *E. coli* pJB37 pFE604T-*ydiE* | - | ED Fig. 2A | Retron over-expression growth assay (trigger validation) |
| p2-*betI* p-empty | *E. coli* pTU175 pFE604T-*betI* | - | ED Fig. 2A | Retron over-expression growth assay (trigger validation) |
| p2-*betI* p-retron | *E. coli* pJB38 pFE604T-*betI* | - | ED Fig. 2A | Retron over-expression growth assay (trigger validation) |
| p2-*betI* p-retron-Δ*rcaT* | *E. coli* pJB51 pFE604T-*betI* | - | ED Fig. 2A | Retron over-expression growth assay (trigger validation) |
| p2-*betI* p-*rcaT* | *E. coli* pJB37 pFE604T-*betI* | - | ED Fig. 2A | Retron over-expression growth assay (trigger validation) |
| p2-*pspG* p-empty | *E. coli* pTU175 pFE604T-*pspG* | - | ED Fig. 2A | Retron over-expression growth assay (trigger validation) |
| p2-*pspG* p-retron | *E. coli* pJB38 pFE604T-*pspG* | - | ED Fig. 2A | Retron over-expression growth assay (trigger validation) |
| p2-*pspG* p-retron-Δ*rcaT* | *E. coli* pJB51 pFE604T-*pspG* | - | ED Fig. 2A | Retron over-expression growth assay (trigger validation) |
| p2-*pspG* p-*rcaT* | *E. coli* pJB37 pFE604T-*pspG* | - | ED Fig. 2A | Retron over-expression growth assay (trigger validation) |
| p1-*pinE* p-empty | *E. coli* pTU175 pNTR-SD-*pinE* | - | ED Fig. 2A | Retron over-expression growth assay (trigger validation) |
| p1-*pinE* p-retron | *E. coli* pJB38 pNTR-SD-*pinE* | - | ED Fig. 2A | Retron over-expression growth assay (trigger validation) |
| p1-*pinE* p-retron-Δ*rcaT* | *E. coli* pJB51 pNTR-SD-*pinE* | - | ED Fig. 2A | Retron over-expression growth assay (trigger validation) |
| p1-*pinE* p-*rcaT* | *E. coli* pJB37 pNTR-SD-*pinE* | - | ED Fig. 2A | Retron over-expression growth assay (trigger validation) |
| p1-empty p-retron-Δ*rcaT* | *E. coli* pJB51 pNTR-SD | - | ED Fig. 2A | Retron over-expression growth assay (trigger validation) |
| p1-empty p-retron | *E. coli* pJB38 pNTR-SD | - | ED Fig. 2A | Retron over-expression growth assay (trigger validation) |
| p2-empty p-retron-Δ*rcaT* | *E. coli* pJB51 pFE604T | - | ED Fig. 2A | Retron over-expression growth assay (trigger validation) |
| p2-empty p-retron | *E. coli* pJB38 pFE604T | - | ED Fig. 2A | Retron over-expression growth assay (trigger validation) |
| p1-*racC* p-empty | *E. coli* pTU175 pNTR-SD-*racC* | - | ED Fig. 4A | RcaT over-expression growth assay (blocker validation) |
| p1-*racC* p-retron | *E. coli* pJB38 pNTR-SD-*racC* | - | ED Fig. 4A | RcaT over-expression growth assay (blocker validation) |
| p1-*racC* p-retron-Δ*rcaT* | *E. coli* pJB51 pNTR-SD-*racC* | - | ED Fig. 4A | RcaT over-expression growth assay (blocker validation) |
| p1-*racC* p-*rcaT* | *E. coli* pJB37 pNTR-SD-*racC* | - | ED Fig. 4A/4B | RcaT over-expression growth assay (blocker validation) |
| p1-*yfbO* p-empty | *E. coli* pTU175 pNT3-*yfbO* | - | ED Fig. 4A | RcaT over-expression growth assay (blocker validation) |

**Supplementary Table 3.** Genotypes of bacterial strains used in this study (3/3).

| | | | | |
|---|---|---|---|---|
| p1-*yfbO* p-retron | *E. coli* pJB38 pNT3-*yfbO* | - | ED Fig. 4A | RcaT over-expression growth assay (blocker validation) |
| p1-*yfbO* p-retron-Δ*rcaT* | *E. coli* pJB51 pNT3-*yfbO* | - | ED Fig. 4A | RcaT over-expression growth assay (blocker validation) |
| p1-*yfbO* p-*rcaT* | *E. coli* pJB37 pNT3-*yfbO* | - | ED Fig. 4A/4B | RcaT over-expression growth assay (blocker validation) |
| p1-*yfbN* p-empty | *E. coli* pTU175 pNT3-*yfbN* | - | ED Fig. 4A | RcaT over-expression growth assay (blocker validation) |
| p1-*yfbN* p-retron | *E. coli* pJB38 pNT3-*yfbN* | - | ED Fig. 4A | RcaT over-expression growth assay (blocker validation) |
| p1-*yfbN* p-retron-Δ*rcaT* | *E. coli* pJB51 pNT3-*yfbN* | - | ED Fig. 4A | RcaT over-expression growth assay (blocker validation) |
| p1-*yfbN* p-*rcaT* | *E. coli* pJB37 pNT3-*yfbN* | - | ED Fig. 4A/4B | RcaT over-expression growth assay (blocker validation) |
| p1-*rtcB* p-empty | *E. coli* pTU175 pNT3-*rtcB* | - | ED Fig. 4A | RcaT over-expression growth assay (blocker validation) |
| p1-*rtcB* p-retron | *E. coli* pJB38 pNT3-*rtcB* | - | ED Fig. 4A | RcaT over-expression growth assay (blocker validation) |
| p1-*rtcB* p-retron-Δ*rcaT* | *E. coli* pJB51 pNT3-*rtcB* | - | ED Fig. 4A | RcaT over-expression growth assay (blocker validation) |
| p1-*rtcB* p-*rcaT* | *E. coli* pJB37 pNT3-*rtcB* | - | ED Fig. 4A/4B | RcaT over-expression growth assay (blocker validation) |
| p1-*mhpR* p-empty | *E. coli* pTU175 pNTR-SD-*mhpR* | - | ED Fig. 4A | RcaT over-expression growth assay (blocker validation) |
| p1-*mhpR* p-retron | *E. coli* pJB38 pNTR-SD-*mhpR* | - | ED Fig. 4A | RcaT over-expression growth assay (blocker validation) |
| p1-*mhpR* p-retron-Δ*rcaT* | *E. coli* pJB51 pNTR-SD-*mhpR* | - | ED Fig. 4A | RcaT over-expression growth assay (blocker validation) |
| p1-*mhpR* p-*rcaT* | *E. coli* pJB37 pNTR-SD-*mhpR* | - | ED Fig. 4A/4B | RcaT over-expression growth assay (blocker validation) |
| p1-*ybeD* p-empty | *E. coli* pTU175 pNTR-SD-*ybeD* | - | ED Fig. 4A | RcaT over-expression growth assay (blocker validation) |
| p1-*ybeD* p-retron | *E. coli* pJB38 pNTR-SD-*ybeD* | - | ED Fig. 4A | RcaT over-expression growth assay (blocker validation) |
| p1-*ybeD* p-retron-Δ*rcaT* | *E. coli* pJB51 pNTR-SD-*ybeD* | - | ED Fig. 4A | RcaT over-expression growth assay (blocker validation) |
| p1-*ybeD* p-*rcaT* | *E. coli* pJB37 pNTR-SD-*ybeD* | - | ED Fig. 4A/4B | RcaT over-expression growth assay (blocker validation) |
| p1-*clpX* p-empty | *E. coli* pTU175 pNT3-*clpX* | - | ED Fig. 4A | RcaT over-expression growth assay (blocker validation) |
| p1-*clpX* p-retron | *E. coli* pJB38 pNT3-*clpX* | - | ED Fig. 4A | RcaT over-expression growth assay (blocker validation) |
| p1-*clpX* p-retron-Δ*rcaT* | *E. coli* pJB51 pNT3-*clpX* | - | ED Fig. 4A | RcaT over-expression growth assay (blocker validation) |
| p1-*clpX* p-*rcaT* | *E. coli* pJB37 pNT3-*clpX* | - | ED Fig. 4A/4B | RcaT over-expression growth assay (blocker validation) |
| p1-*ydaW* p-empty | *E. coli* pTU175 pNT3-*ydaW* | - | ED Fig. 4A | RcaT over-expression growth assay (blocker validation) |
| p1-*ydaW* p-retron | *E. coli* pJB38 pNT3-*ydaW* | - | ED Fig. 4A | RcaT over-expression growth assay (blocker validation) |
| p1-*ydaW* p-retron-Δ*rcaT* | *E. coli* pJB51 pNT3-*ydaW* | - | ED Fig. 4A | RcaT over-expression growth assay (blocker validation) |
| p1-*ydaW* p-*rcaT* | *E. coli* pJB37 pNT3-*ydaW* | - | ED Fig. 4A/4B | RcaT over-expression growth assay (blocker validation) |
| p1-*chpS* p-empty | *E. coli* pTU175 pNT3-*chpS* | - | ED Fig. 4A | RcaT over-expression growth assay (blocker validation) |
| p1-*chpS* p-retron | *E. coli* pJB38 pNT3-*chpS* | - | ED Fig. 4A | RcaT over-expression growth assay (blocker validation) |
| p1-*chpS* p-retron-Δ*rcaT* | *E. coli* pJB51 pNT3-*chpS* | - | ED Fig. 4A | RcaT over-expression growth assay (blocker validation) |
| p1-*chpS* p-*rcaT* | *E. coli* pJB37 pNT3-*chpS* | - | ED Fig. 4A/4B | RcaT over-expression growth assay (blocker validation) |
| p1-empty p-*rcaT* | *E. coli* pJB37 pNTR-SD | - | ED Fig. 4B | RcaT over-expression growth assay (blocker validation) |
| p2-empty p-*rcaT* | *E. coli* pJB37 pFE604T | - | ED Fig. 4B | RcaT over-expression growth assay (blocker validation) |
| p2-*dicC* p-*rcaT* | *E. coli* pJB37 pFE604T-*dicC* | - | ED Fig. 4B | RcaT over-expression growth assay (blocker validation) |
| p2-*clpA* p-*rcaT* | *E. coli* pJB37 pFE604T-*clpA* | - | ED Fig. 4B | RcaT over-expression growth assay (blocker validation) |
| p1-empty p-retron-Δ*rcaT* | *E. coli* pNTR-SD pJB51 | NT32113 | ED Fig. 5A | Retron over-expression growth assay |
| p1-empty p-retron | *E. coli* pNTR-SD pJB38 | NT32167 | ED Fig. 5A | Retron over-expression growth assay |
| p1-$dam^{Ec}$ p-retron-Δ*rcaT* | *E. coli* pJB81 pJB51 | NT16885 | ED Fig. 5A | Retron over-expression growth assay |
| p1-$dam^{Ec}$ p-retron | *E. coli* pJB81 pJB38 | NT16884 | ED Fig. 5A | Retron over-expression growth assay |
| p1-$dam^{S\ Tm}$ p-retron-Δ*rcaT* | *E. coli* pJB83 pJB51 | NT16907 | ED Fig. 5A | Retron over-expression growth assay |
| p1-$dam^{S\ Tm}$ p-retron | *E. coli* pJB83 pJB38 | NT16906 | ED Fig. 5A | Retron over-expression growth assay |
| p1-$dam^{P1}$ p-retron-Δ*rcaT* | *E. coli* pJB85 pJB51 | NT16913 | ED Fig. 5A | Retron over-expression growth assay |
| p1-$dam^{P1}$ p-retron | *E. coli* pJB85 pJB38 | NT16912 | ED Fig. 5A | Retron over-expression growth assay |
| Δ*xseA* p1-empty p-$retron^{WT}$ | *E. coli* Δ*xseA*::kan pNTR-SD pJB38 | NT32168 | ED Fig. 5B | Retron over-expression growth assay |
| Δ*xseA* p1-*dam* p-$retron^{WT}$ | *E. coli* Δ*xseA*::kan pNTR-SD-*dam* pJB38 | NT32170 | ED Fig. 5B | Retron over-expression growth assay |
| Δ*xseA* p1-empty p-$retron^{mut}$ | *E. coli* Δ*xseA*::kan pNTR-SD pJB128 | NT32172 | ED Fig. 5B | Retron over-expression growth assay |
| Δ*xseA* p1-*dam* p-$retron^{mut}$ | *E. coli* Δ*xseA*::kan pNTR-SD-*dam* pJB128 | NT32174 | ED Fig. 5B | Retron over-expression growth assay |
| Δ*xseA* p1-empty p-empty | *E. coli* Δ*xseA*::kan pNTR-SD pTU175 | NT32164 | ED Fig. 5B | Retron over-expression growth assay |
| Δ*xseA* p1-*dam* p-empty | *E. coli* Δ*xseA*::kan pNTR-SD-*dam* pTU175 | NT32166 | ED Fig. 5B | Retron over-expression growth assay |
| p-retron | *S* Tm Δretron::FRT Δ*araBAD*::kan pJB38 | NT32177 | ED Fig. 5C | msDNA isolation |
| p-$retron^{mut}$ | *S* Tm Δretron::FRT Δ*araBAD*::kan pJB128 | NT32178 | ED Fig. 5C | msDNA isolation |
| p-retron-ΔrcaT-Eco9 p1-recE | *E. coli* pJB95 pNTR-SD-*recE* | - | ED Fig. 6A | Retron-Eco9 over-expression growth assay |
| p-retron-Eco9 p1-recE | *E. coli* pJB118 pNTR-SD-*recE* | - | ED Fig. 6A | Retron-Eco9 over-expression growth assay |
| p-retron-ΔrcaT-Eco9 p1-empty | *E. coli* pJB95 pNTR-SD | - | ED Fig. 6A | Retron-Eco9 over-expression growth assay |
| p-retron-Eco9 p1-empty | *E. coli* pJB118 pNTR-SD | - | ED Fig. 6A | Retron-Eco9 over-expression growth assay |
| WT p-retron-Δ*rcaT* | *S* Tm Δretron::FRT Δ*araBAD*::kan pJB51 | NT16658 | ED Fig. 6B | msDNA isolation |
| Δ*xseA* p-retron-Δ*rcaT* | *S* Tm Δ*xseA*::FRT Δ*araBAD*::kan pJB51 | NT16659 | ED Fig. 6B | msDNA isolation |
| p-rcaT-Eco9 p1-empty | *E. coli* pJB117 pNTR-SD | - | ED Fig. 6C | RcaT-Eco9 over-expression growth assay |
| p-rcaT-Eco9 p1-racC | *E. coli* pJB117 pNTR-SD-*racC* | - | ED Fig. 6C | RcaT-Eco9 over-expression growth assay |
| p2-RT-Eco1 WT | *S* Tm pFE604T-B21_00839 | - | ED Fig. 7 | Cold-sensitivity growth assay |
| p2-RT-Eco1 Δ*rrtT* | *S* Tm Δ*rrtT*::FRT pFE604T-B21_00839 | - | ED Fig. 7 | Cold-sensitivity growth assay |
| p2-RT-Eco1 Δretron | *S* Tm Δretron::FRT pFE604T-B21_00839 | - | ED Fig. 7 | Cold-sensitivity growth assay |
| p2-RT-Eco1 Δ*msrmsd* | *S* Tm Δ*msrmsd*::FRT pFE604T-B21_00839 | - | ED Fig. 7 | Cold-sensitivity growth assay |
