## Supplementary Table 4. Description of plasmids used in this study. for "Phage proteins block and trigger retron toxin/antitoxin systems"

**Supplementary Table 4.** Description of plasmids used in this study (1/1).

| **Plasmid code** | **Plasmid name** | **Backbone** | **Used for** | **Antibiotic resistance** | **Reference** |
|---|---|---|---|---|---|
| pNTR-SD | pNTR-SD | pNT3 | Over-expression of *E. coli* genes. | Ampicillin | 25 |
| pFE604T | pFE604T | pFE604T | Over-expression of *E. coli* genes. | Tetracycline/Gentamycin | 26 |
| pJB37 | pTU175-pBAD-*rcaT* | pTU175 | Over-expression of *rcaT* . | Spectinomycin | 24 |
| pJB38 | pTU175-pBAD-retron | pTU175 | Over-expression of retron. | Spectinomycin | 24 |
| pJB51 | pTU175-*msrmsd*-*rrtT* | pTU175 | Over-expression of *msrmsd*-*rrtT* . | Spectinomycin | 24 |
| pJB81 | pNTR-SD-$dam^{Ec}$ | pNTR-SD | Over-expression of *dam* from *E. coli*. | Ampicillin | This work |
| pJB83 | pNTR-SD-$dam^{STm}$ | pNTR-SD | Over-expression of *dam* from *S* Tm. | Ampicillin | This work |
| pJB85 | pNTR-SD-$dam^{P1}$ | pNTR-SD | Over-expression of *dmt* (dam-homologue) from Enterobacteria Phage P1. | Ampicillin | This work |
| pJB91 | pBAD33-rcaT(Eco9) | pBAD33 | Over-expression of rcaT(Eco9). | Chloramphenicol | 24 |
| pJB92 | pBAD33-retron(Eco9) | pBAD33 | Over-expression of retron(Eco9). | Chloramphenicol | 24 |
| pJB93 | pTU175-pBAD-msrmsd(Eco9)-RT(Sen2) | pTU175 | Construction of pJB95. | Spectinomycin | This work |
| pJB95 | pTU175-pBAD-msrmsd(Eco9)-RT(Eco9) | pTU175 | Over-expression of msrmsd(Eco9)-RT(Eco9). | Spectinomycin | This work |
| pJB117 | pTU175-pBAD-rcaT(Eco9) | pTU175 | Over-expression of rcaT-Eco9. | Spectinomycin | This work |
| pJB118 | pTU175-pBAD-retron(Eco9) | pTU175 | Over-expression of retron-Eco9. | Spectinomycin | This work |
| pJB128 | pTU175-pBAD-retron:*msd*-[GATC→GTTC] | pTU175 | Over-expression of retron carrying a GATC → GTTC mutation in *msd* . | Spectinomycin | This work |
| pKM1 | pNTR-SD-$msrmsd^{WT}$ | pNTR-SD | Over-expression of *msrmsd* . | Ampicillin | 24 |
| pKM3 | pNTR-SD-$msrmsd^{mut}$ | pNTR-SD | Over-expression of *msrmsd* carrying a G→T mutation in the branching G of the *msr* region. | Ampicillin | This work |
| pTU175 | pTU175-empty | pTU175 | Cloning. | Spectinomycin | 75 |
