## Supplementary Table 5. Description of construction of plasmids used in this study. for "Phage proteins block and trigger retron toxin/antitoxin systems"

**Supplementary Table 5.** Description of construction of plasmids used in this study (1/1).

| Plasmid code | Plasmid construction comments | Vector | Insert |
|---|---|---|---|
| pJB81 | Insert was amplified using primers JB328 and JB330 from strain *E. coli* , and ligated in HindIII-SalI cut-pNTR-SD. | pNTR-SD | HindIII-*dam* $^{Ec}$ -SalI |
| pJB83 | Insert was amplified using primers JB341 & JB342 from strain *S* Tm, and ligated in HindIII-SalI cut-pNTR-SD. | pNTR-SD | HindIII-*dam* $^{S\ Tm}$-SalI |
| pJB85 | Insert was amplified using primers JB339 & JB340 from a P1 phage lysate (from strain *E. coli* ), and ligated in HindIII-SalI cut-pNTR-SD. | pNTR-SD | HindIII-*dam* $^{P1}$-SalI |
| pJB93 | Insert was amplified using primers JB370 & JB390 from plasmid pJB118, and ligated in XmaI-XbaI cut-pJB51. | pJB51 | XmaI-msrmsd(Eco9)-XbaI |
| pJB95 | Insert was amplified using primers JB391 & JB392 from plasmid pJB118, and ligated in XbaI-HindIII cut-pJB93. | pJB93 | XbaI-RT(Eco9)-HindIII |
| pJB117 | Insert was amplified using primers JB368 & JB416 from plasmid pJB91, and ligated in XmaI-HindIII cut-pJB38. | pJB38 | XmaI-rcaT(Eco9)-HindIII |
| pJB118 | Insert was amplified using primers JB370 & JB417 from plasmid pJB92, and ligated in XmaI-HindIII cut-pJB38. | pJB38 | XmaI-msrmsd-rcaT-RT(Eco9)-HindIII |
| pJB128 | Q5-site directed mutagenesis on plasmid pJB38. Mutagenic primers JB433 & JB434 were used to mutate the 5'-GATC-3' duplex on the msDNA towards 5'-GTTC-3'. | pJB38 | - |
| pKM3 | Insert was amplified using primers KM_R_02 & KM_R_03 from plasmid pJB38, and ligated in HindIII-SalI cut-pNTR-SD. | pNTR-SD | HindIII-msrmsd$^{mut}$-SalI |
