## Supplementary Table 6. List of primers used in this study. for "Phage proteins block and trigger retron toxin/antitoxin systems"

**Supplementary Table 6.** List of primers used in this study (1/1).

| **Primer code** | **Primer name** | **Primer sequence** | **Primer description** | **Used in** |
|---|---|---|---|---|
| JB328 | dam-Ec_HindIII_For | TTCATCATAAGCTT atgAAGAAAAATCGCGCTTTTTTGAAG | Green: anneals to start of *dam* gene (*E. coli*), blue: restriction site, black: restriction enzyme buffer-sequence. | pJB81 plasmid construction. |
| JB330 | dam-Ec_SalI_Rev | TTCATCATGTCGAC ttaTTTTTTCGCGGGTGAAACGACTC | Green: anneals to end of *dam* gene (*E. coli*), blue: restriction site, black: restriction enzyme buffer-sequence. | pJB81 plasmid construction. |
| JB341 | dam-STm_HindIII_For | TTCATCATAAGCTT atgAAAAAAAATCGCGCTTTTTTGAAGTGG | Green: anneals to start of *dam* gene (*S*Tm), blue: restriction site, black: restriction enzyme buffer-sequence. | pJB83 plasmid construction. |
| JB342 | dam-STm_SalI_Rev | TTCATCATGTCGAC ttaTTTTCTTGCAGGCGTTGCG | Green: anneals to end of *dam* gene (*S*Tm), blue: restriction site, black: restriction enzyme buffer-sequence. | pJB83 plasmid construction. |
| JB339 | dam-P1_HindIII_For | TTCATCATAAGCTT atgAAAGAGCTGTGCTATGGATCTG | Green: anneals to start of dmt gene (phage P1), blue: restriction site, black: restriction enzyme buffer-sequence. | pJB85 plasmid construction. |
| JB340 | dam-P1_SalI_Rev | TTCATCATGTCGAC ttaGAGCAATATACCCAAAACGTCATGAGC | Green: anneals to end of dmt gene (phage P1), blue: restriction site, black: restriction enzyme buffer-sequence. | pJB85 plasmid construction. |
| JB368 | rcaT-Eco9_XmaI_For | TTCATCATCCCGGG atgAGTGATTCTGCATATATGAATGCATTAC | Green: anneals to start of rcaT-Eco9 gene, blue: restriction site, black: restriction enzyme buffer-sequence. | pJB117 plasmid construction. |
| JB370 | msr-Eco9_XmaI_For | TTCATCATCCCGGG ATCTCGCATCTTTTGCTATCCTAACC | Green: anneals to start of msr-Eco9 in plasmid pJB118, blue: restriction site, black: restriction enzyme buffer sequence. | pJB93, pJB118 plasmid construction. |
| JB390 | msd-Eco9_XbaI_Rev | TTCATCATTCTAGA CCACCACAAACAAAAATAATAGGAGGCAT | Green: anneals to +107 nts of rcaT-Eco9 in plasmid pJB118, blue: restriction site, black: restriction enzyme buffer sequence. | pJB93 plasmid construction. |
| JB391 | RT-Eco9_XbaI_For | TTCATCATTCTAGA atgGAGTTGCTAAGTTATATTTCAAAAGGTC | Green: anneals to start of rt-Eco9 in plasmid pJB118, blue: restriction site, black: restriction enzyme buffer sequence. | pJB95 plasmid construction. |
| JB392 | RT-Eco9_HindIII_Rev | TTCATCATAAGCTT ttaAATTTCATATTTTGAAATTTCGCTGATTACATCATCC | Green: anneals to end of rt-Eco9 in plasmid pJB118, blue: restriction site, black: restriction enzyme buffer sequence. | pJB95 plasmid construction. |
| JB416 | rcaT-Eco9_HindIII_Rev | TTCATCATAAGCTT CcatTctaAGCCCCTCCA | Green: anneals to end of rcaT-Eco9 gene, blue: restriction site, black: restriction enzyme buffer-sequence. | pJB117 plasmid construction. |
| JB417 | RT-Eco9_HindIII_Rev | TTCATCATAAGCTT ttaAATTTCATATTTTGAAATTTCGCTGATTACATCATCC | Green: anneals to end of RT-Eco9 gene, blue: restriction site, black: restriction enzyme buffer-sequence. | pJB118 plasmid construction. |
| JB433 | msd-GATC→GTTC_For | AAGTCTCTAGGAACGGTAGAAAACTTTCTAGCGCCTC | Green: anneals to end of *msd* region, red: T→A mutation of GATC motif. | pJB128 plasmid construction. |
| JB434 | msd-GATC→GTTC_Rev | CTAGGAACGGTAGAAAACTTTCTAGCGCAACCCTAAAAG | Green: anneals to start of *msd* region, red: T→A mutation of GATC motif. | pJB128 plasmid construction. |
| KM_R_02 | msrmsd$^{mut}$_For | CAAGCAAGCTT ACATCACTCTTTATCGTTAGG | Green: anneals to start of *msr* region, red: G→T mutation of branching G, blue: restriction site, black: restriction enzyme buffer sequence | pKM3 plasmid construction. |
| KM_R_03 | msrmsd$^{mut}$_Rev | GAGCTGTCGAC CAAACATTGATTTAAACGTTATG | Green: anneals to start of *rcaT* gene, blue: restriction site, black: restriction enzyme buffer sequence | pKM3 plasmid construction. |
